# Gut Microbial Impact on Colitis and Colitis-Associated Carcinogenesis in a Primary Sclerosing Cholangitis-IBD Model

**DOI:** 10.1101/2024.10.03.616279

**Authors:** Muyiwa Awoniyi, Billy Ngo, Vik Meadows, Deniz Coskuner, Stephanie A. Montgomery, Morgan Farmer, Bo Liu, Huiping Zhou, Jeffery Roach, Thaddeus Stappenbeck, R. Balfour Sartor

## Abstract

**Background and Aims:** Primary sclerosing cholangitis (PSC) associated inflammatory bowel diseases (IBD) increase colorectal dysplasia and malignancy risk. Current mouse models do not adequately replicate human PSC-IBD, limiting mechanistic understanding and therapeutic development. This study uses *Mdr2/Il10* double knockout (DKO) mice to examine microbiota roles in mediating colitis, colitis-associated colorectal dysplasia and hepatobiliary inflammation/fibrosis.

**Goal:** Develop and phenotype a chronic spontaneous PSC-IBD mouse model, emphasizing colitis, colonic dysplasia, hepatobiliary inflammation/ fibrosis and the functional roles of resident microbiota.

**Methods:** We utilized germ-free (GF) and specific-pathogen-free *Mdr2/Il10* DKO, *Il10*^-/-^ and *Mdr2*^-/-^ mice to model PSC-IBD. We monitored colonic dysplasia progression, colitis kinetics and severity by lipocalin-2, cytokine measurement, and tissue evaluations of colon and liver. We manipulated the microbiome to assess its functional effects.

**Results:** DKO mice exhibited age- and region-specific accelerated colitis and spontaneous colonic dysplasia progressing to high-grade invasive adenocarcinomas. Despite aggressive colonic inflammation, DKO mice showed reduced hepatic fibrosis, increased hepatic reparative macrophages, and matrix metalloproteinase activity compared to *Mdr2*^-/-^ mice. GF DKO had heightened liver inflammation and mortality with absent colitis and colonic dysplasia, reversed with microbial reconstitution from DKO mice. Changes in DKO primary/secondary bile acid profiles mirrored those in PSC-IBD.

**Conclusion:** The *Mdr2/Il10* DKO model mirrors key factors in PSC-IBD patients in terms of inflammation and carcinogenesis. We found an important role for the dysbiotic microbiota in DKO mice for disease onset and progression. Targeting microbiota and bile acid metabolism may provide promising strategies for developing effective PSC-IBD therapies.

## INTRODUCTION

Primary sclerosing cholangitis (PSC) is a chronic cholestatic liver disease characterized by peribiliary inflammation progressing to hepatic fibrosis. PSC leads to liver failure in nearly 1/6 of affected subjects ^1^. No current medical therapies exist beyond liver transplant, which is not curative ^2^. Up to 80% of PSC patients exhibit concurrent inflammatory bowel disease (IBD), while PSC occurs in 7.5-15.1% of IBD patients^2–4^. While the PSC-IBD nexus is underscored by a distinct clinical and molecular signature, the underlying pathophysiological mechanisms remain elusive ^5–9^.

PSC patients develop characteristic endoscopic ileocolonic inflammation, with right colonic predominance, relative rectal sparing, and ’backwash’ ileitis ^3,4^. Patients with PSC-ulcerative colitis (UC) represent over 85% of the PSC IBD subset and have a 4-10-fold greater risk for developing colorectal neoplasia compared to those with UC alone ^10–12^. Emerging evidence reveals a discordance between mild endoscopic findings and more active histological and molecular inflammation ^13–15^. This suggests occult subclinical inflammation with a significant risk of malignancy^14, 16^. These patients also experience earlier onset of neoplasia, accounting for approximately 8% of PSC-related mortality ^3, 11^ and are more susceptible to pouchitis post-colectomy, often necessitating early antibiotic intervention, suggesting the microbiome’s role in disease pathogenesis^1,3,17^.

Current animal models fail to encapsulate the multifaceted pathophysiology and clinical manifestations of human PSC-IBD ^18^. The *Mdr2*^-/-^mouse, a commonly employed PSC model, exhibits defective biliary phosphatidylcholine transport, leading to liver pathology reminiscent of human PSC ^19^. The mice do not, however, develop spontaneous colonic inflammation characteristic of PSC-IBD ^20^. To induce this colonic inflammation, chemically induced models have been used to dissect liver-gut axis interactions and explore the interactions of experimental colitis and hepatic inflammation and fibrosis. For example, transient oral administration of dextran sulfate sodium (DSS) to *Mdr2* deficient mice induces mixed effects on colonic inflammation, increased intestinal permeability, but with reduced hepatobiliary injury. ^21, 22, 23^ However, the DSS model bears notable limitations, such as variable colonic disease contingent on microbial composition, primarily distal colonic involvement, and non-T and B cell dependent colitis, limiting its translational relevance ^24, 25^. Alternatively, researchers have used *Il10*^-/-^ mice as a colitis model in combination with a hepatotoxin to mimic the chronicity and T-cell-dependence seen in colitis patients^26^. IL-10 deficiency leads to spontaneous colitis in the presence of resident microbiota but does not induce hepatobiliary inflammation ^27^. Applying hepatotoxic agents like intraperitoneal administered carbon tetrachloride (CCl_4_) to *Il10*^-/-^ mice provokes severe hepatic fibrosis, impaired hepatocyte proliferation and elevated TNF-α levels^28, 29^. However, the chemically induced hepatic inflammation in this latter model is hepatocyte directed injury, not biliary, therefore not faithfully representing the biliary-focused inflammation of PSC. Thus, neither model fully captures the complexity of spontaneous right-sided predominant colitis and dysplasia, and the chronic progression of inflammation and fibrosis observed in human PSC-IBD ^30^.

To investigate the complex pathogenesis of PSC-IBD, we crossbred *Mdr2*^-/-^ with *Il10*^-/-^ mice to generate the *Mdr2*/*Il10* DKO line on the C57Bl/6 background. The amalgamation of these gene deletions synergistically models the hepatic and colonic phenotypes observed in PSC-IBD patients. Bedke et al. recently demonstrated a similar mouse model as an experimental PSC-IBD model, but with divergent findings and varying approaches^11^.

Our model exhibits age and genotype specific changes in host microbiota associated with spontaneous early-onset colitis and accelerated dysplasia in the right colon with attenuated hepatic inflammation and fibrosis. Our analyses reveal a distinctive microbial functional profile in the DKO model, suggesting an interactive role between bacterial components, bile acid metabolism, and disease progression. DKO mice have early onset, aggressively progressive right-sided colorectal dysplasia and cancer, driven by accelerated colitis suggesting a link between biliary inflammation and colitis-associated colorectal oncogenesis. Absent microbiota enhances liver-mediated mortality but prevents colitis in DKO mice. Microbial transfer induces colitis but conveys hepatoprotection, underscoring the differential influences of intestinal microbiota on colitis and liver inflammation. Furthermore, co-housing DKO with *Il10*^-/-^ mice attenuated colitis and colorectal carcinogenesis supporting a strong role for microbial mitigation of inflammation and associated dysplasia. These insights not only enrich our understanding of PSC-IBD pathogenesis but also signal a promising avenue for microbial-targeted therapies.

## MATERIAL AND METHODS

### SPF and Germ-Free Double Knockout Mice

Specific-pathogen-free (SPF) and germ-free (GF) *Mdr2/Il10* double knockout (DKO), *Il10*^-/-^, *Mdr2^+/-^*/*Il10*^-/-^*, Mdr2*^-/-^, and wild-type (WT) C57BL/6 mice were generated and weaned at 3-4 weeks. Mice were aged 6-45 weeks. GF mice were generated by Cesarean sections and bred at the National Gnotobiotic Rodent Resource Center at UNC-Chapel Hill. *Mdr2^+/-^/Il10*^-/-^ were bred to each other to obtain *Mdr2*^-/-^ and *Il10*^-/-^ littermates. See Supplementary Materials and Methods for detailed information.

### Assessment of Colitis

Severity of colitis was evaluated by serial fecal lipocalin-2 ELISA and blinded histologic scoring as previously described ^31^. See Supplementary Materials and Methods for detailed procedures.

### Animal Interventional Experiments

SPF mice were treated with antibiotics or underwent fecal content transplantation. See Supplementary Materials and Methods for detailed protocols.

### Histology

Tissue sections were stained and examined for histopathological features. Neoplastic lesions were scored by small animal pathologist. See Supplementary Materials and Methods for detailed procedures.

### Co-housing DKO and *Il10*^-/-^ Model

DKO and *Il10*^-/-^ mice were co- or singly-housed, treated with Azoxymethane (AOM) or PBS, and necropsied at 4 months post-treatment. See Supplementary Materials and Methods for detailed protocols.

### Immunohistochemical Staining and Quantification of Hepatic Macrophage Markers

Liver tissue sections were stained for macrophage markers, scanned, and analyzed for positive staining. See Supplementary Materials and Methods for detailed protocols.

### Statistical Analysis

Statistical significance was assessed using ANOVA and post-hoc Student’s *t*-test. The Kruskal-Wallis test, PCoA, PERMANOVA, and ALDEx2 were used for microbial diversity and differential abundance analyses. All p-values adjusted for multiple comparison using Benjamini–Hochberg procedure.

### Ethics Statement/Animal Husbandry

Animal studies were conducted in accordance with NIH guidelines. All experiments were approved by the IACUC at UNC (Protocol no. 18-266) and Cleveland Clinic (Protocol no. 2281). See Supplementary Materials and Methods for details.

## RESULTS

### Colonic tumor progression in DKO Mice

*Mdr2*^-/-^*/Il10*^-/-^ (double knock out, DKO) mice developed spontaneous features of intestinal tumorigenesis with initial dysplasia followed by invasive adenocarcinoma. At 16 weeks of age, 50% of DKO mice showed high grade epithelial dysplasia (SF1A), particularly in the right colon (Fig. 1A-B) while dysplasia in littermate control mice (*Il10*^-/-^), ranged from mild to absent (Fig. 1A-D). Older DKO mice (22-44 weeks) showed moderate to high-grade epithelial dysplasia throughout the colon, while dysplasia in *Il10*^-/-^ mice ranged from absent to moderate (Fig. 1C, D). In 9-month-old mice, we observed colonic invasive adenocarcinoma in 50% of DKO mice; in contrast, none of the *Il10*^-/-^ mice developed colonic adenocarcinomas (Fig. 1E). The DKO colons showed classic histologic features of moderately differentiated invasive mucinous adenocarcinoma with submucosal infiltration of irregular colonic epithelial glandular structures that were associated with pools of PAS+ mucin (Fig 1F-G, S1C). Thus, DKO mice may be leveraged to interrogate mechanisms in the gut-liver axis underlying colorectal cancer progression.

**Figure 1:**
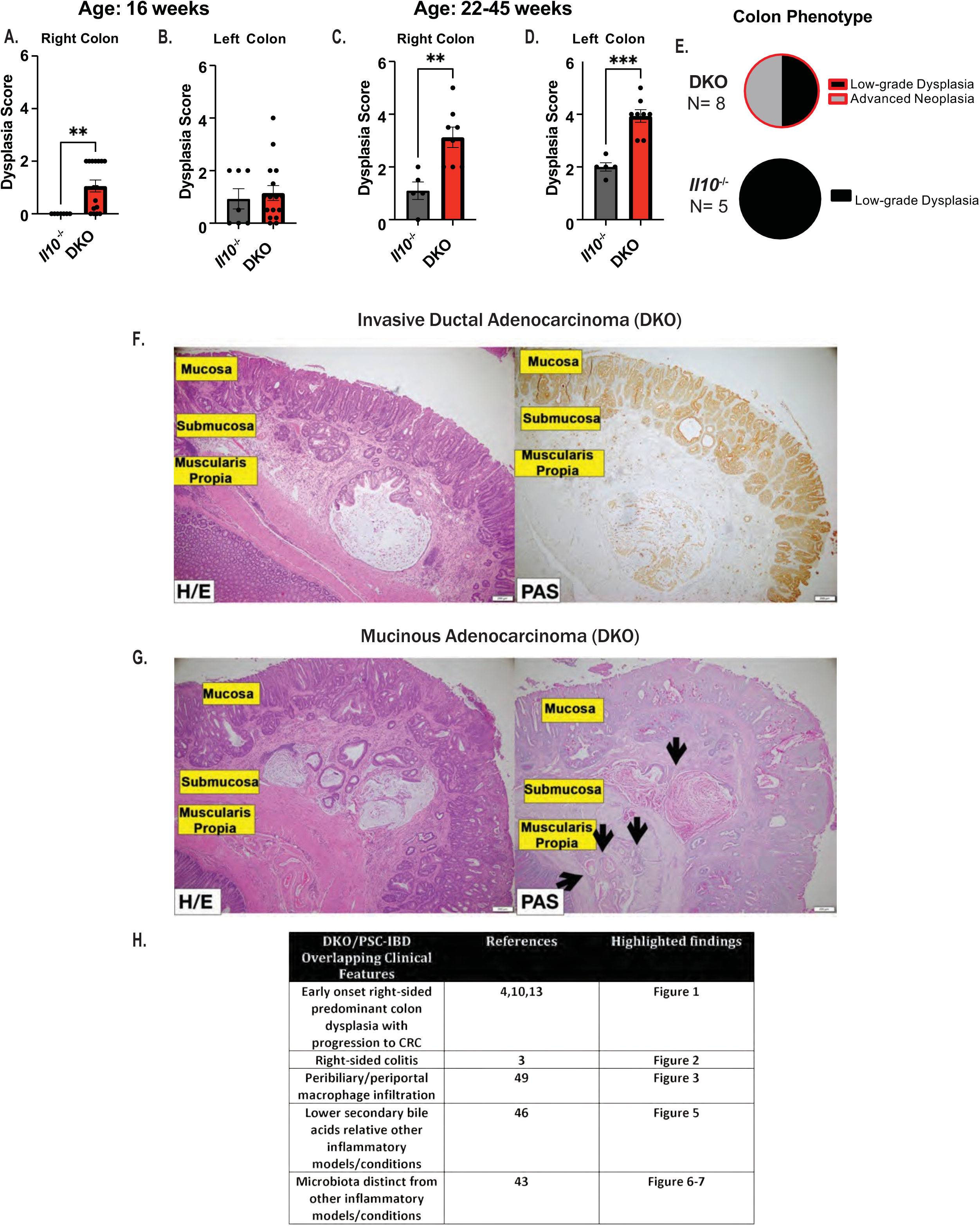
PSC-IBD Model Mice Exhibit Amplified Dysplasia. (A-B) Regional dysplasia scores in 16-week-old DKO versus *Il10*^-/-^ mice show differential severity; n = 16 for DKO and n = 7 for *Il10*^-/-^. (C-D) Dysplasia scoring of older mice (22–45 weeks) of for right (cecum and proximal colon) and left (distal colon) regions; n = 8 for DKO and n = 5 for *Il10*^-/-^. (E) Pie chart summarizing the relative severity of dysplasia in 22–45-week-old DKO compared to *Il10*^-/-^ mice. Representative images (40x magnification) of distal colon from a 42-week-old DKO mouse. Left panel: Hematoxylin and Eosin (H&E) staining; Right panel: pan-cytokeratin (panCK) staining demonstrating epithelial cell origin. (G) Sample photomicrographs of colonic mucinous adenocarcinoma at 40X magnification in the proximal colon of another 42-week-old DKO mouse. Left panel: H&E staining; Right panel: Periodic acid-Schiff (PAS) staining with arrows highlighting invasive dysplastic glands. Colonic mucosal layers (in yellow). (H) Summary of key overlapping human and murine model findings. Data are presented as mean ± SEM. Statistical significance was assessed via ANOVA for multiple group comparisons and Student’s t-test for pairwise analyses, with *P<0.05, **P<0.01, ***P<0.001, ****P<0.0001 indicating increasing levels of significance.

### Early-life Colitis as a Precursor to Colorectal Dysplasia in DKO Mice

We hypothesize that adenocarcinomas in older DKO mice arose in part from enhanced early-life colonic inflammation. We found elevated fecal lipocalin-2 (Lcn2) levels as early as 4 weeks of age in DKO mice but not control mice (littermate *Il10*^-/-^ and *Mdr2*^-/-^) (Fig. 2A, SF2B). Notably, by 6 weeks of age, DKO colons (both right and left) and ilea exhibited higher levels of inflammation as compared to *Il10*^-/-^ mice, (higher histopathological scores measuring immune infiltration (Fig. 2B, C, D, G; SF2L) or qPCR of colonic *iNos2*mRNAs (SF2C)). This finding was independent of mouse weights (SF2A). Interestingly, at 16 weeks, colonic H+E sections of DKO and *Il10*^-/-^ mice showed similar amounts of inflammation (Fig. 2A, E, F, SF2M). However, 16-week-old DKO mice maintained elevated colonic *Tnfa* and *iNos2* RNA expression as compared to WT mice (Fig SF2J-K). Colitis and mouse weights were not different in *Mdr^+/-^*/*Il10*^-/-^ and *Il10*^-/-^ littermate control mice (SFN-O). Thus, the appearance of early onset colonic inflammation in DKO mice suggests that there exists a critical period of colitis-driven susceptibility to dysplasia.

**Figure 2:**
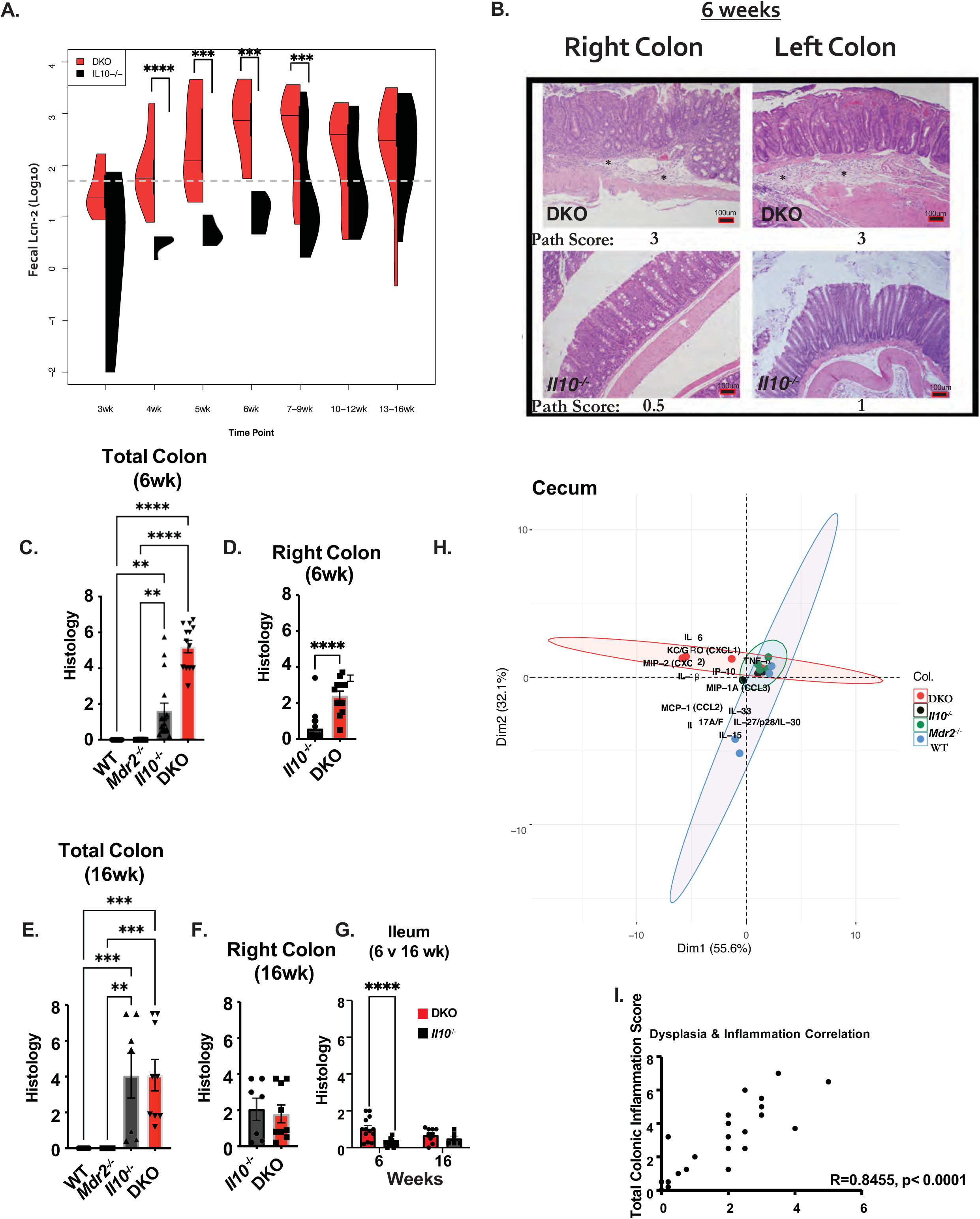
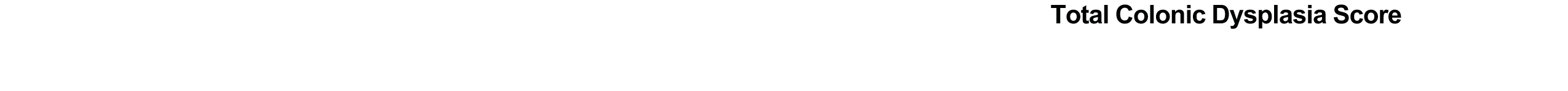
Early Inflammation in DKO Mice Sets the Stage for Colorectal Dysplasia. (A) Longitudinal tracking of fecal lipocalin-2 (Lcn-2) in SPF DKO, *Il10*^-/-^, *Mdr2^−/−^*, and wild-type WT mice, shown as a split violin plot. The dashed line represents the threshold below which Lcn-2 levels are clinically insignificant, as confirmed by histological validation. Sample sizes range from n=5 to n=23 for DKO and *Il10*^-/-^ . (B) Histological sections of proximal and distal colonic tissue from 6-week-old DKO and *Il10*^-/-^ mice stained with H&E (C, E) Compiled blinded pathology scores for total colonic inflammation (cecum + ascending + distal colon) and (D, F) right colon inflammation (cecum and ascending colon) in DKO, *Mdr2*^-/-^, and WT mice at ages 6 and 16 weeks. (G) Ileal inflammation scores for the compared groups at both time points. (H) Principal coordinates analysis (PCoA) displaying the variations in cytokine and chemokine profiles in cecal tissue among genotypes: DKO, *Mdr2*^-/-^, *Il10*^-/-^, and WT (n=4/group). Proteins analyzed include IL-1β, MCP-1 (CCL2), MIP1 (CCL3), MIP2 (CXCL2), TNF-α, KC/Gro (CXCL1), IL-6, IL-33, IL-17a/f, IP-10, IL-15, and IL-27/p28/IL-30. (I) Correlation coefficient (R = 0.8455). PERMANOVA p-value = 0.0158. Pairwise comparison adjusted p-values: DKO vs. WT q = 0.0313; DKO vs. *Il10*^-/-^ q=0.2857; DKO vs. *Mdr2*^-/-^ q=0.1095; WT vs. *Il10*^-/-^ q=0.1714; WT vs. *Mdr2*^-/-^ q=0.12; *Il10*^-/-^ vs. *Mdr2*^-/-^ q=0.5143. Statistical analyses included ANOVA for group comparisons and Student’s t-test for individual contrasts.

Peak differences in fecal Lcn2 levels between DKO and *Il10*^-/-^ mice in early life coincided with higher levels of cecal inflammatory cytokine and chemokine expression (Fig. 2H, SF1D-E). DKO mice showed enriched cecal interferon gamma (IFN-g)-induced Protein 10 (IP-10 also referred to as CXCL10) (SF1E). The principal coordinate analysis (PCA) of cytokine and chemokine expression further differentiated the DKO phenotype from single knockout and WT mice, underscoring a distinct inflammatory signature inherent to this model (Fig. 2H). Correlation analyses confirmed a robust positive association between total colonic inflammation and dysplasia (Fig. 2I), supporting our hypothesis that early colonic inflammation in DKO mice drives dysplasia onset.

### Decreased Portal Inflammation and Liver Fibrosis in DKO mice is Associated with Increased Reparative Macrophage Infiltration

We next evaluated the relationship of hepatic inflammation, injury and fibrosis in DKO. We hypothesized that heightened colitis in DKO mice would worsen hepatobiliary injury and fibrosis. Paradoxically we found that DKO mice had reduced periportal inflammation and liver injury relative to *Mdr2^-/-^* mice (Fig. 3A-C), associated with similar reductions in IL-17a and IL-15 (Fig. 3D-E). The kinetics of ALT levels over time differed between *Mdr2*^-/-^ and DKO. *Mdr2*^-/-^showed earlier and ultimately higher levels of ALT expression while DKOs showed delayed progression (Fig. 3C). Thus, liver and colons in DKO mice show a temporal mismatch of inflammation, with early onset colitis, but reduced and delayed hepatic inflammation.

**Figure 3:**
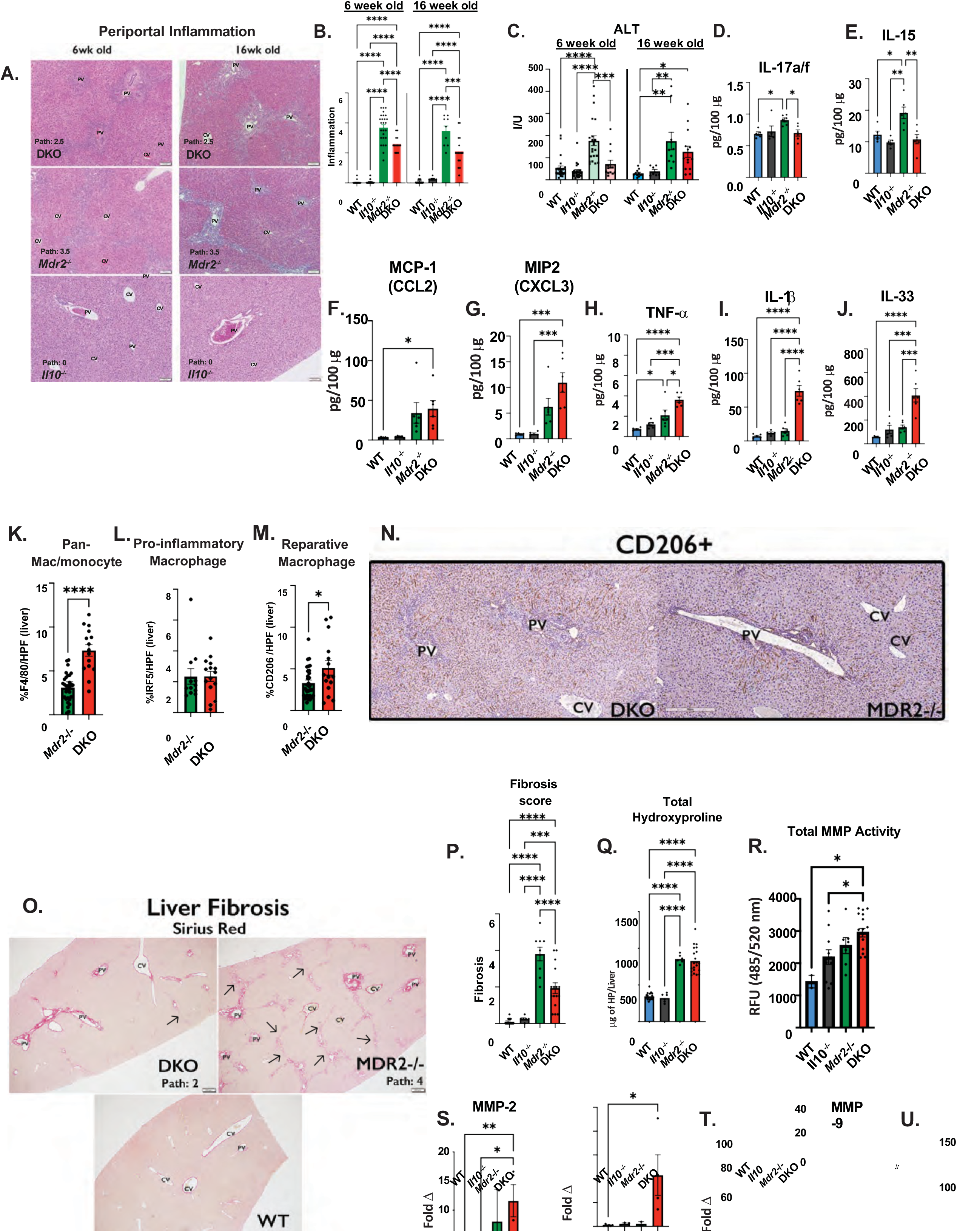
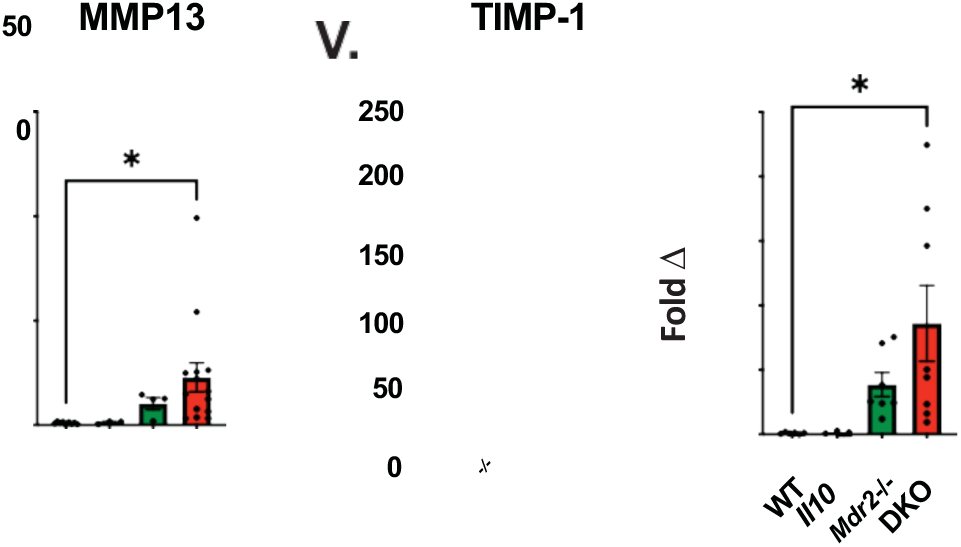
Reduced Periportal Inflammation and Fibrosis with Increased Infiltration of Reparative Macrophages in PSC-IBD Mouse Model. (A) Sample photomicrograph of H/E liver staining (100X), (B) composite scoring of periportal inflammation and (C) ALT at 6 and 16 weeks. (D-J) Histogram of hepatic IL-17a/f, IL-15, CCL2/MCP-1, CCL3/MIP2, TNF-α, IL-1β and IL-33 production by multiplex panel (n= 5-6/group). (K-M) Quantification of hepatic macrophage populations via immunohistochemistry, employing markers F4/80 for pan-macrophages, IRF-5 for pro-inflammatory macrophages, and CD206 for reparative macrophages n=DKO (n=13) and *Mdr2^−/−^* (n=29). (N) Representative immunohistochemical photomicrograph at 200X magnification showing CD206+ cells in hepatic tissue of DKO and *Mdr2^−/−^* mice. (O-P) Sample photomicrograph of Sirius Red staining for liver with magnification at 100X, and total liver fibrosis scoring. (Q) Total liver hydroxyproline content (reported as μg hydroxyproline per whole liver, normalized to liver weight). (R-V) Total liver matrix-metalloproteinase (MMP) enzymatic activity, and liver RNA expression of MMP-2, -9, and -13, along with TIMP-1. Data are represented as mean ± SEM. Statistical analyses utilized ANOVA for group comparisons and Student’s t-test for pairwise contrasts.

While most pro-inflammatory cytokines showed lower levels of expression in DKO livers, a subset of cytokines and chemokines (Fig. 3F-J), were elevated; notably IL-33, which induces differentiation of reparative macrophages that contribute to the resolution of inflammation and tissue repair was elevated in DKO mice ^32^. In addition, DKO mice showed higher numbers of F4/80+ myeloid cells in the liver relative to *Mdr2*^-/-^ mice (Fig. 3K) as well as enriched expression of CD206, a marker of reparative macrophages (Fig. 3M-N). Markers for inflammatory macrophages did not differ between these two groups. These results support a model of reparative inflammation in DKO livers, that occurs concurrently with the development of a pro-inflammatory colitis.

This model is further supported by DKO livers exhibiting less histologic fibrosis evidenced by Sirius Red staining (Fig. O-P) despite similar levels of accumulated hydroxyproline content at 16 weeks of age (Fig. 3Q). Examination of specific molecules that control extracellular matrix dynamics including matrix metalloproteinases (MMPs) and tissue inhibitor of matrix metalloproteinase (TIMP)-1 showed enhanced MMP protease activity and expression in DKO liver tissue relative to non-liver disease controls (Fig. 3R-V). The increased reparative macrophage phenotype and elevated levels of these matrix modifiers suggest a mechanism of the reduced fibrotic phenotype of DKO mice.

### The Intestinal Microbiome Safeguards against Hepatic Inflammation and Fibrosis while Driving Colitis

We hypothesized that the microbiota protects against hepatic inflammation and fibrosis while inducing colitis in DKO mice. To support this hypothesis, we found that germ-free (GF) *Mdr2*^-/-^ and DKO mice exhibited reduced survival compared to *Il10*^-/-^ or *Mdr2^+/-^*/*Il10*^-/-^ littermate controls (Fig. 4A). Pairwise comparison showed GF *Mdr2*^-/-^ had accelerated mortality relative to DKO mice (SF3G), associated with heightened liver histological and biochemical fibrosis, ductular reaction (Fig. 4H-I, SF3R) but lacked colitis (Fig. 4D-F). These results demonstrate the hepatoprotective role of resident microbiota.

**Figure 4:**
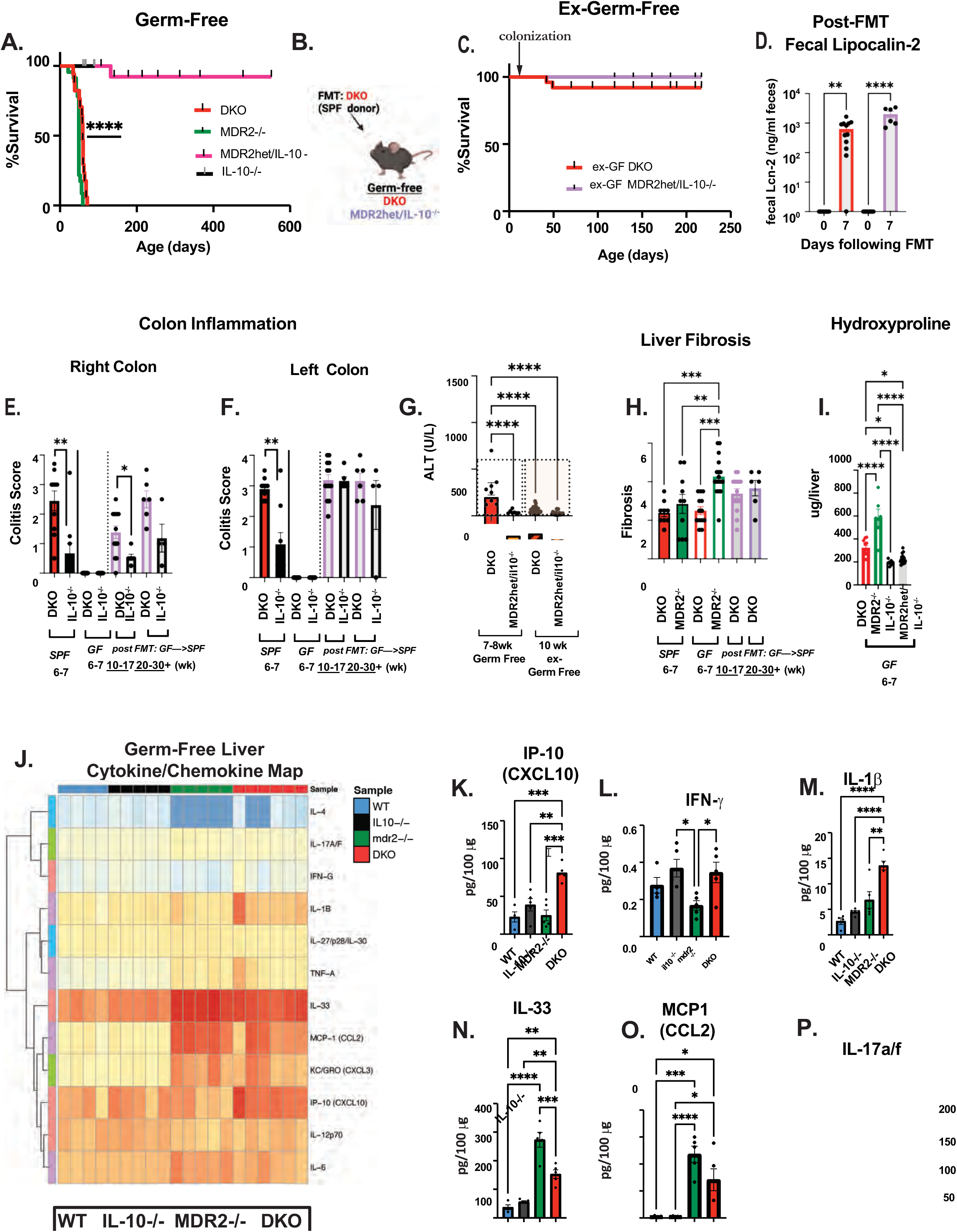
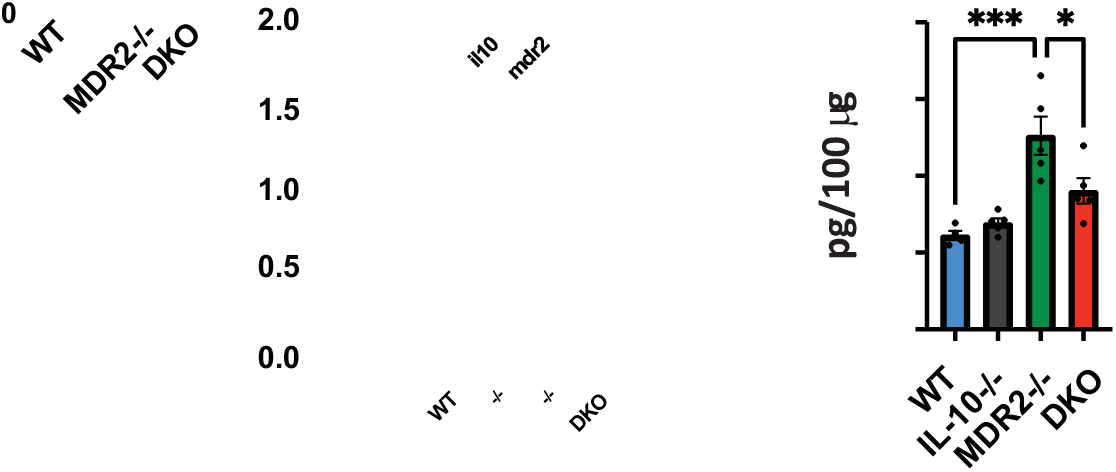
Microbial Reconstitution Through Fecal Transplant Boosts Survival and Mitigates Liver Dysfunction While Provoking Colitis in GF DKO Mice. (A) Survival curves using Kaplan-Meier estimates for germ-free (GF) DKO mice relative to *Mdr22*^-/-^, *Mdr22*het/ *Il10*^-/-^, and *Il10*^-/-^ strains and (B-C) reconstituted “ex-GF” DKO and *Mdr2*het/ *Il10*^-/-^ from DKO pooled donor. n=5–6 per group. (D) Fecal Lcn-2 pre/7-day post oral fecal transfer (5-9/group). (E-F) Histological segmental colitis and (H) liver fibrosis score of age matched SPF vs GF and longitudinal assess of reconstituted GF DKO and *Il10*^-/-^ mice, n=6 -13 at each group. (G.) Serum ALT levels before and after FMT of DKO vs *Mdr2*het/ *Il10*^-/-^littermate controls (n=10-15/group). (I) Total hepatic hydroxyproline content (in μg HYP/whole liver) across various GF mouse genotypes. (J) Heatmap of differentially expressed multiplex liver cytokines/chemokines in GF DKO mice. (K-P) Histogram representation of multiplex liver cytokine profile: IP-10/CXCL10, IFN-g, IL-1*β*, IL-33, MCP-1/CCL2 and IL-17a/f. Sample sizes were n=5–6 per group. All results are presented as mean ± SEM. Survival rates were analyzed with the Log-rank (Mantel-Cox) test, while ANOVA and Student’s t-test were utilized for additional group comparisons.

Microbial reconstitution from DKO specific pathogen free (SPF) donors, restored survival to DKO mice, attenuated liver injury (Fig. 4G, SF3O) and liver fibrosis progression (like *Mdr2*^-/-^ mice), but with rapid induction of colitis (unlike *Mdr2*^-/-^ mice) (Fig. 4D-F). Like GF outcomes, histologic analysis revealed that antibiotic exposure increased liver inflammation and fibrosis in DKO and *Mdr2^-/-^* mice (SF4A-B). In contrast, antibiotics attenuated colitis in SPF DKO and *Il10*^-/-^ mice demonstrated by reduced fecal Lcn-2 and histologic inflammation (SF4C-D). These findings underscore the pivotal role of microbiota in driving colonic disease.

The cytokine/chemokine protein multiplex in GF DKO and *Mdr2*^-/-^ mouse livers showed greater levels of CXCL10 (IP-10), IFN-γ, and IL-1β (Fig. 4J-M) and less IL-33, CCL2, CCL3, and IL-17a/f in DKO compared to *Mdr2*^-/-^ mice (Fig. 4N-P, SF3H). Under GF conditions, the increased monocyte/macrophages and reparative macrophages observed in DKO relative to *Mdr2*^-/-^ livers in SPF conditions were no longer evident (SF3P, S-T).

In summary, our findings illustrate the divergent impact of the microbiota on hepatic and colonic pathologies in DKO mice, offering potential avenues for microbiota-centric mechanisms.

### Distinct Cecal Bile Acid Composition in DKO Mice Indicates Altered Metabolic Pathways

We hypothesized that DKO mice exhibit distinct cecal bile acid compositions due to altered microbial bile acids metabolism. Recognizing the pivotal role of bile acids as functional microbial metabolites and their association with the pathogenesis of PSC, we probed bile acid metabolites in this model. Only *Mdr2^-/-^* had significantly reduced total cecal bile acids (TBA) compared to WT (Fig. 5B) while DKO and *Mdr2^-/-^* had lower secondary bile acids compared to *Il10^-/-^* mice (Figs. 5C-E). This reduction could signify deficient microbial bile acid 7-*α*-dehydroxylase activity.

**Figure 5:**
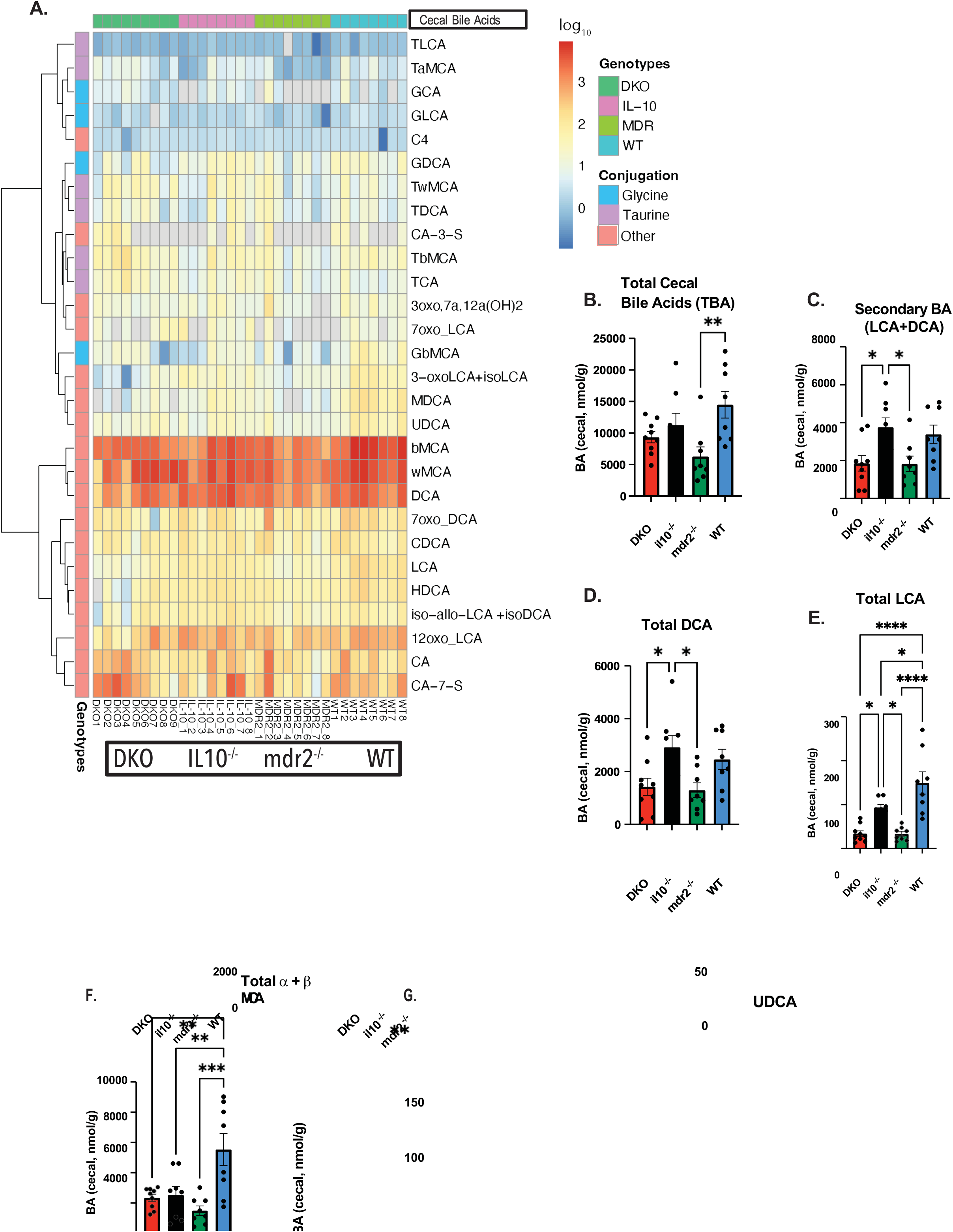
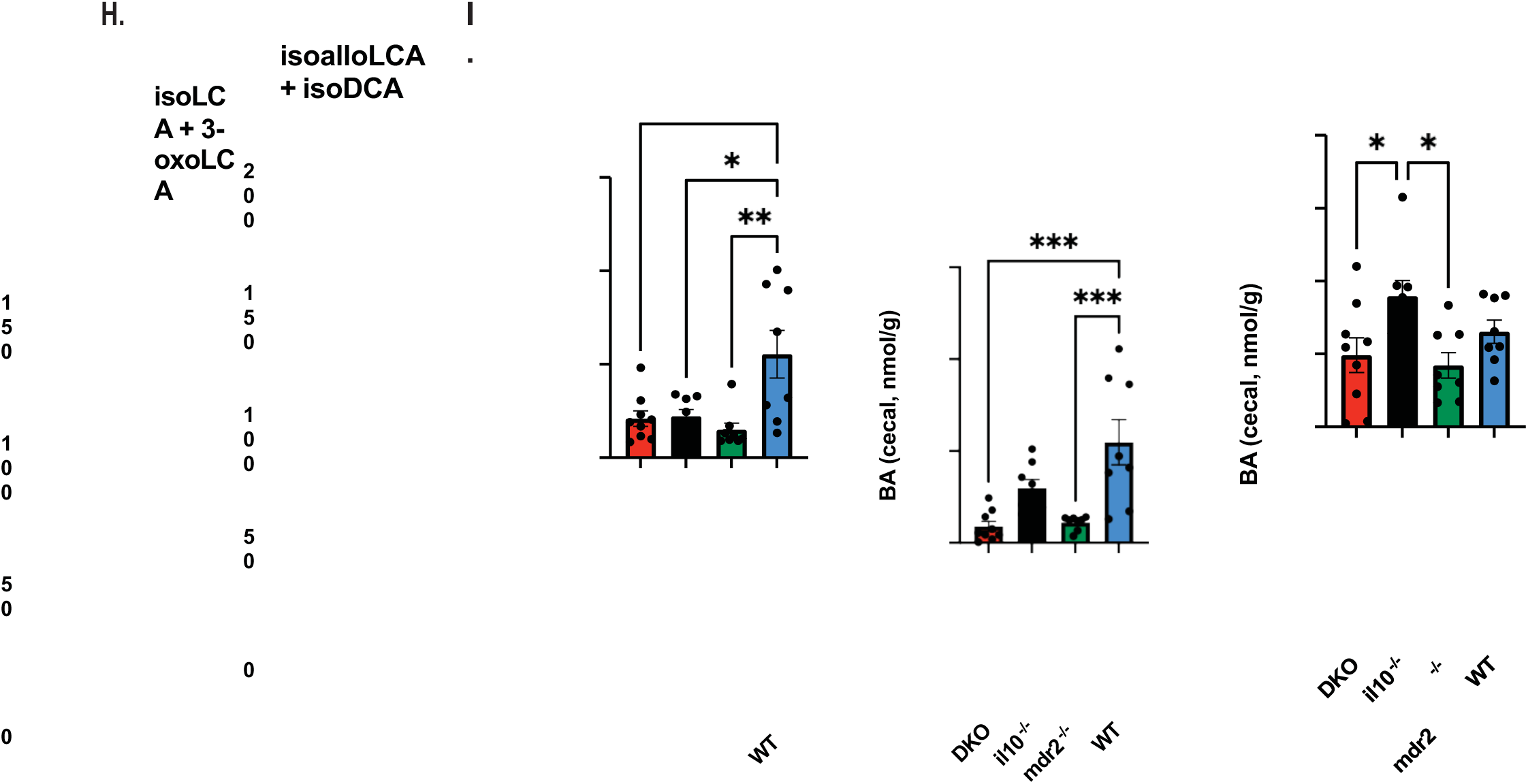
Distinct Bile Acid Signatures in DKO Mice Reflect *Mdr2*-Dependent Metabolic Alterations. (A) The log-transformed heatmap displays the varied spectrum of cecal bile acids across four genotypes— DKO, *Il10*^-/-^, *Mdr2*^-/-^, and WT—at 6-8 weeks. (B) Histogram analysis of TBA across genotypes. (C–E) Comparative quantification of secondary bile acids, with specific focus on total secondary bile acids (LCA+DCA), DCA, and LCA, to illustrate genotype-specific metabolic transformations. (F) Differential analysis of α- and MCA, highlighting variances in primary bile acid profiles among genotypes. (G) Differential analysis of UDCA. (H) Differential analysis of RORγt-inhibitory bile acids isoallolithocholic (isoLCA) and 3-oxoLCA. (I) Differential analysis of FOXP3-inducing bile acids (isoLCA and isoDCA). Sample sizes are n=7–8 per genotype, with results presented as mean ± SEM. Statistical evaluations employed ANOVA for group comparisons and Student’s t-test for pairwise assessments. Significance is noted by p-value thresholds set within the study.

Furthermore, we noted diminished levels of muricholic acids (MCAs) and ursodeoxycholic acid (UDCA) in DKO, *Mdr2^-/-^*, and *Il10^-/-^* mice compared to WT mice (Figs. 5F-G). Inhibitors of Th17 differentiation, isolithocolic acid (isoLCA) and 3-oxoLCA ^33, 34^ were substantially lower in DKO and *Mdr2*^-/-^ mice (Fig. 5H) yet without corresponding changes in *foxp3* cecal tissue mRNA expression across genotypes (SF5K). Interestingly, we found unaltered cecal RNA expression of *RORγt*, a critical Th17 and inducible Treg transcription factor (SF5J). No significant differences occurred in the *fxr-fgf15* (ileum)-*cyp7α1* (liver) axis across the mouse models (SF5L-N), indicating an alternative mechanism driving the observed bile acid profile. We subsequently investigated the gut microbiota’s contribution to these metabolic changes and their subsequent impact on the pathophysiology of PSC-IBD.

### Differential Microbial Abundance Highlights Unique Interactions in DKO Mice

We hypothesized that specific intestinal microbial populations in DKO mice differentially influence disease phenotypes in the livers and colon. To evaluate this, we performed whole genome sequencing on fecal samples from DKO and control mice. Analysis revealed no significant variations in alpha diversity (Fig. 6A). A PCoA plot based on Bray-Curtis dissimilarity showed that DKO mice harbor a distinct microbial signature (Fig. 6B).Genotype pairwise comparison of the relative differential abundance highlighted higher abundance of various species in DKO, *Mdr2*^-/-^ and *Il10*^-/-^ mice. In particular, *Faecalibaculum rodentium* was more abundant in the DKO and *Mdr2*^-/-^ mice vs *Il10*^-/-^ mice (Fig. 6C, blue). *F. rodentium* has an anti-tumorigenic effect in colorectal cancer models via short-chain fatty acid (SCFA)-mediated inhibition of NFATc3 proliferative activity ^35^. At 8 weeks, DKO mice had higher levels of fecal *F. rodentium* DNA compared to *Il10*^-/-^ mice. (SF6A), however, we did not find any baseline differences in cecal SCFA at 6-8 weeks of age vs other groups (SF6B-D).

**Figure 6:**
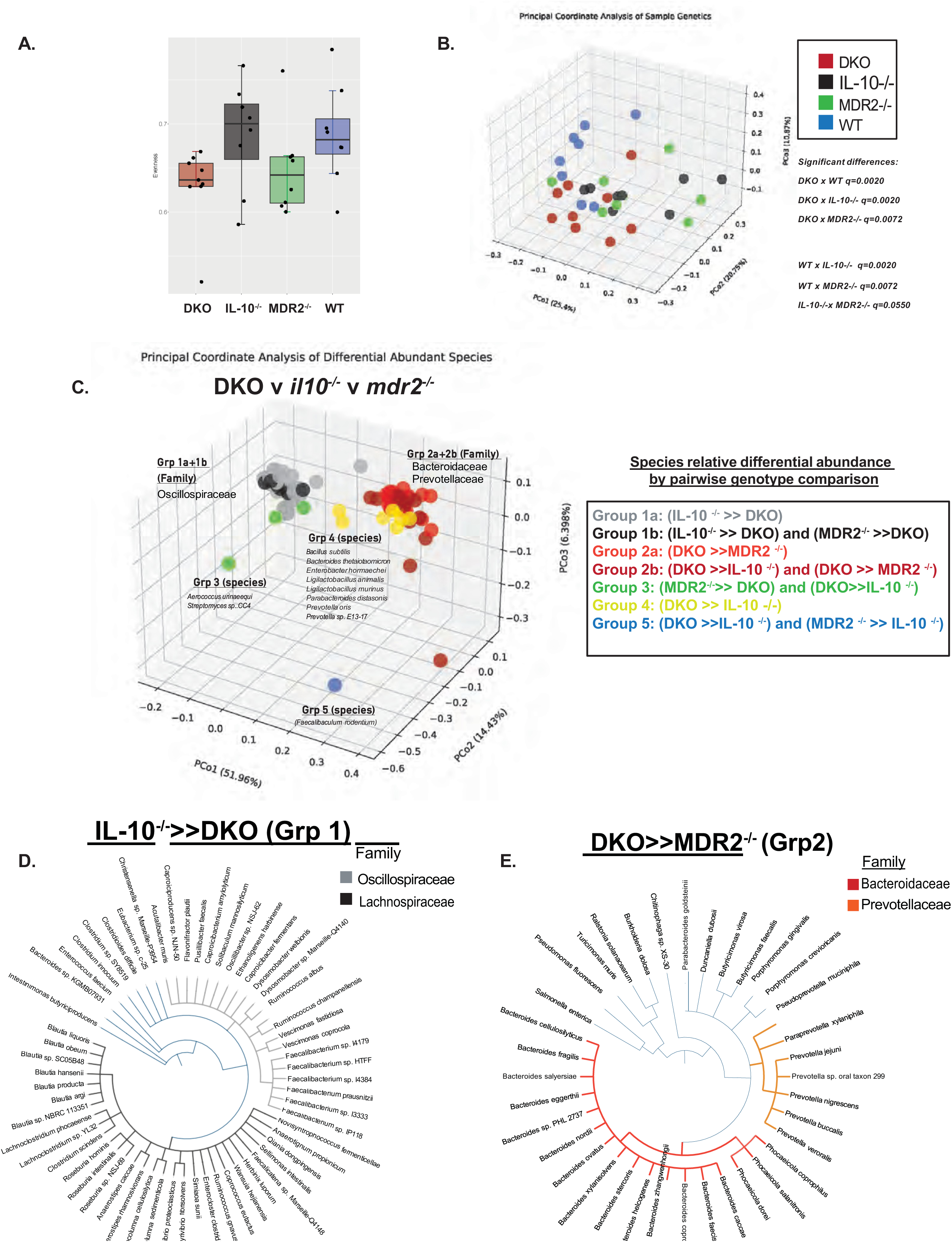
DKO Mice Harbor a Distinctive Bacterial Taxonomic Profile at Baseline. (A) α-diversity of bacterial communities is presented through Pielou’s Evenness index box plots of 6-week-old DKO and respective controls. The overall Kruskal-Wallis p-value is 0.0723, with no pairwise comparisons reaching significance below an adjusted p-value of 0.1. (B) A 3D PCoA plot delineates the genetic influences on microbial composition, utilizing Bray-Curtis dissimilarity. Individual mouse samples are depicted as points, color-coded by genotype, with PERMANOVA p-value 0.001. Adjusted p-values from genotype pairwise comparisons are detailed numerically. (C) Microbial species with differential abundance identified in any pairwise genotype comparisons are illustrated. A color-coding system categorizes species based on differential abundance patterns between genotypes: gray for species differentially more relatively abundant in *Il10*^-/-^ versus DKO, black for species differentially more relatively abundant in *Mdr2*^-/-^ and *Il10*^-/-^ compared to DKO, salmon for species differentially more relatively abundant in DKO versus *Mdr2*^-/-^, dark red for species differentially more relatively abundant in DKO versus *Il10*^-/-^ and DKO versus *Mdr2*^-/-^, green for species differentially more relatively abundant in *Mdr2*^-/-^ versus DKO and also differentially more relatively abundant in DKO versus *Il10*^-/-^, yellow for species differentially more relatively abundant in DKO versus *Il10*^-/-^, and blue specifically for *Faecalibaculum rodentium’s* differential abundance where DKO and *Mdr2*^-/-^ are differentially more abundant versus *Il10*^-/-^. (D–E) Cladograms illustrate taxa in *Il10*^-/-^ versus DKO (Group 1) and DKO versus *Mdr2*^-/-^ (Group 2) respectively. Taxonomic families Lachnospiraceae, Bacteroidaceae, Prevotellaceae, and Oscillospiraceae are highlighted in corresponding colors. (N= 8 mice/group).

Cladogram analysis showed distinct microbial composition of differentially abundance families in different groups. There was an overrepresentation of Bacteroidaceae and Prevotellaceae in DKO versus *Mdr2*^-/-^ mice (Group 2) as well as overrepresentation of Lachnospiraceae and Oscillospiraceae families in *Il10*^-/-^ versus DKO mice (Group 1; Fig. 6D-E).

### Attenuation of Colitis and Colorectal Dysplasia Through Intergenotypic Horizontal Transfer

We tested the bidirectional potential of horizontal microbial exchange between genetic strains of mice. At weaning, we separated genotype-specific strains showing microbiome divergent composition described in Fig. 6. We used this experimental scheme to test two models: co-housing adult mice would transmit colitogenic bacteria from DKO to *Il10*^-/-^ mice, or conversely, transfer protective microbial constituents from *Il10*^-/-^ to DKO mice. Our co-housing experiments, starting at age of 6–8-week-old mice, evaluated the influence of the microbiota on colonic phenotype, colorectal dysplasia and liver pathologies. To accelerate the experimental timeline of dysplasia in DKO mice, we used azoxymethane (AOM).

Our results indicated that singly housed DKO mice exhibited diminished weight gain over four months post-AOM treatment, whereas co-housed DKO mice weight changes were aligned with *Il10^-/-^* from this weight loss, indicating a stabilizing effect of the exchanged microbiota (Fig. 7B). Microbial exchange by co-housing decreased total colonic histologic inflammatory scores in DKO mice, as reflected by attenuated right-sided colitis, with minimal changes in distal colonic scores (Fig. 7C). *Tnf*α proximal colonic mRNA expression was attenuated in co-housed DKO mice, although distal colonic histology and TNFα protein levels remained unchanged (Fig. 7D-E).

**Figure 7:**
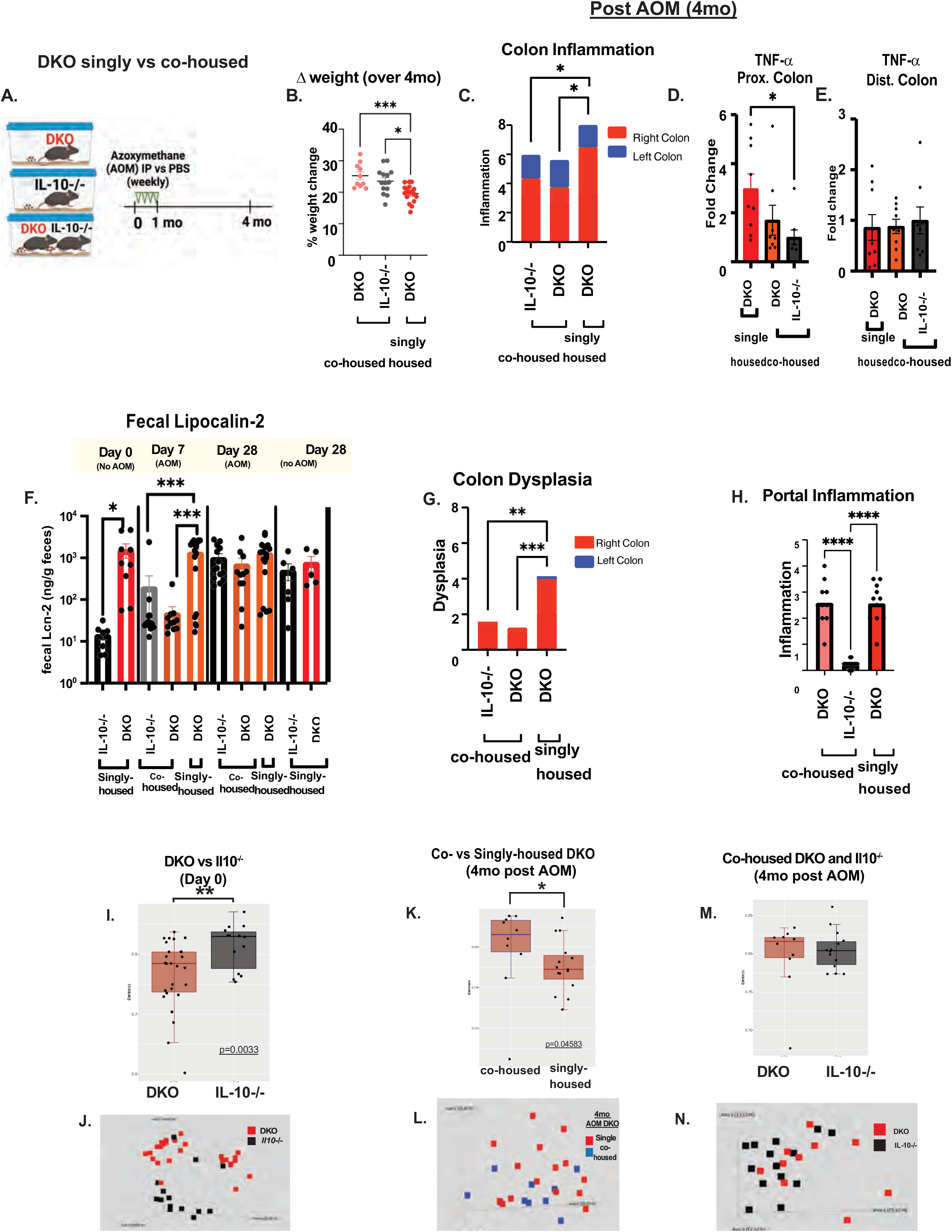
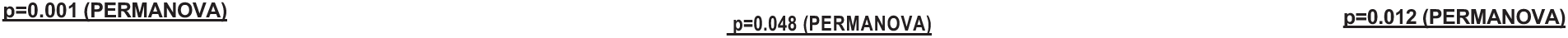
Co-Housing with *Il10*^-/-^ Mice Modulates Microbial Profile and Reduces AOM-Induced Dysplasia in DKO Mice. (A) Experimental co-housing design of singly- vs co-housed SPF DKO with *Il10*^-/-^ mice starting at 8-week-old exposed to AOM or phosphate-buffered saline (PBS) treatments over four months. (B) Longitudinal assessment of (B) *Δ* weight change in singly vs co-housed DKO mice. Single vs co-housed DKO and *Il10*^-/-^ post treatment assessment of (C) colonic inflammation score, (D-E) cecal segmental colonic RNA expression, (F) fecal Lcn2, (G) colonic dysplasia score, and (H) portal inflammation from singly DKO vs cohoused *Il10^-/-^* and DKO mice. (I-N) 16S Microbiota diversity analysis through Pielou’s Evenness and PCoA of beta diversity for DKO and *Il10*^-/-^ mice under various conditions: pre-treatment housing, post-AOM treatment co-housing, and over the experimental timeline. n=10-20 mice per group.

Longitudinal tracking of fecal Lcn2 levels across DKO and *Il10*^-/-^ mice showed that co-housing altered colitis kinetics. Initial Lcn2 values at 8 weeks in singly housed SPF DKO mice were significantly higher than in *Il10*^-/-^ mice (Fig. 7F). Co-housing rapidly reduced Lcn2 levels in DKO mice to levels similar to *Il10*^-/-^ cagemates by day 7; (Fig. 7F). By day 28, Lcn2 levels in co-housed *Il10*^-/-^ mice were similar to DKO mice, suggesting equilibrated inflammatory responses due to microbial sharing. These kinetics demonstrate a protective role of transferred *Il10*^-/-^ mouse microbiota to DKO mice in early phases of colitis.

Protective *Il10*^-/-^ microbiota influences were also apparent in the reduced right colonic dysplastic changes observed in co-housed DKO mice, highlighting the microbiome’s role in tempering colitis-associated dysplasia (Fig. 7G). No discernible differences in portal inflammation were evident between singly or co-housed DKO mice, indicating that microbial transfer from a non-liver inflammatory mouse did not influence DKO liver disease (Fig. 7H).

### Microbiome Dynamics Post-AOM Treatment and Co-Housing

We tested the hypothesis that co-housing DKO mice with *Il10*^-/-^ mice would lead to a stable microbial shift associated with altered colonic inflammation and tumorigenesis by performing fecal 16S rRNA gene sequencing analysis. Alpha diversity was higher in *Il10*^-/-^ mice compared to DKO mice before AOM treatment and co-housing (Fig. 7I). These mice were littermates separated at weaning. PCoA of beta diversity values before co-housing showed genotype-specific microbial clustering, indicating divergent microbial populations (Figs. 6B, 7J). After four months of co-housing, the alpha diversity of DKO and *Il10*^-/-^ mice normalized (Fig. 7M), and the beta diversity showed a shift from baseline (Fig. 7J), suggesting successful microbial integration among cagemates. Notably, co-housed DKO had a greater alpha diversity than individually housed DKO mice (Fig. 7K and I). PCoA revealed minimal separation between samples from co-housed versus individually housed DKO mice (Fig. 7L). Co-housed DKO mice (versus singly house DKO mice) showed reduced Bacteroidaceae and Prevotellaceae abundance, with concomitantly increased bile acid and fatty acid-modifying genus Turicibacter ^36^ (SF7F).The genus Faecalibaculum was found in significantly higher relative abundance prior to co-housing (SF7J). *Faecalibaculum rodentium* was confirmed by qPCR prior to cohabitation but waned over time independent of AOM treatment (SF7K).

In AOM-treated *Il10*^-/-^ mice, no differences were noted in alpha diversity with single versus cohousing (SF7I). The PCoA exhibited contraction akin to that of the *Il10^-/-^* mice post-treatment and cohousing (SF7J). Several genera displayed significant shifts in relative abundance for single vs. co-housed mice: Alloprevotella, Parasutterella, and Erysipelatoclostridium were significantly more prevalent after cohousing relative to baseline DKO mice (SF7G). Conversely, genera such as Enterorhabdus, Alistipes, Rikenella, and Clostridia UCG-014 were relatively more abundant prior to treatment and cohousing, indicating a significant microbial shift in response to environmental and treatment conditions (SF7G). Alloprevotella and Erysipelatoclostridium prevalence increased in 4-months cohoused *Il10*^-/-^ mice vs. baseline (SF7J). We did not find presence of previously published genotoxic bacteria (SF Table1) by 16s analysis. These observations illuminate the profound impact of co-housing on the gut microbiome of both DKO and *Il10*^-/-^ mice. The distinct microbial patterns before and after co-housing highlight the dynamic nature of the gut microbiome and underscore the potential for environmental factors to substantial change microbiota.

## DISCUSSION

We crossbred *Mdr2^-/-^* and *Il10^-/-^* mouse strains to create a model of PSC-IBD to study microbial influences on combined hepatobiliary and colonic inflammation. These DKO mice display many features of clinical PSC-IBD (Fig. 1H), most importantly early right-sided colitis, dysplasia and invasive colonic carcinoma. Our results provide new insights into differential organ specific effects as microbial reconstitution studies showed that DKO microbiota induce colitis and liver inflammation but conversely protects against hepatobiliary fibrosis.

Relevance of the DKO model is provided by clinical observations highlighting significant disparities between endoscopic and histologic evaluations of colitis, suggesting histologic examination is more sensitive for detecting subclinical inflammation ^13–15^. Notably, a Norwegian study found a 2.67-fold increase in the rate of colonic inflammation detection by histology versus endoscopy^14^. Additionally, a cohort of PSC-IBD patients who developed colorectal neoplasia, showed higher endoscopic and histologic indications of right colon inflammation prior to dysplasia ^37,38^. Given these data suggesting long-standing subclinical colonic inflammation contribute to higher colitis-associated dysplasia rates in PSC-UC patients compared to UC patients alone, the DKO model provides a valuable system to explore these hypotheses.

A recent study using *Mdr2^-/-^; Il10^-/-^* DKO mice found reduced endoscopic colitis and increased colonic FOXP3+ Tregs compared to *Il10^-/-^* mice, highlighting phenotypic differences in colitogenic activity depending on the microbiota used for engraftment ^39^. Various technical and experimental differences could explain these contrasting findings, with facility-specific microbiomes likely being the most influential factor. This common occurrence^40,41^ sets the stage for future studies to identify region specific colitogenic microbes leading to dysplasia and cancer in PSC-IBD patients.

Follow-up work by the same lab showed increased FOXP3 expression in PSC-IBD patients is associated with pro-inflammatory colonic expression of *IL17A* and *IFNG*, potentially reflecting a compensatory inflammatory response ^15^. Further support comes from association of IL-17+FOXP3+ DP cells with colitis-associated cancers in PSC-IBD patients ^38^ with unique host microbial signatures. The microbiome-mediated colitis in our DKO model precedes colonic carcinogenesis and may explain early subclinical inflammatory events that shape later disease manifestations in PSC patients^4,37,38,42^.

The microbiome of DKO mice exhibits decreased diversity, reflecting host genotype-specific alterations, that mirror differences between PSC-UC and UC patients ^43^. Animal husbandry details are relevant to interpreting our results. *Mdr2^+/-^; Il10^-/-^* were crossed to obtain *Mdr2^-/-^; Il10^-/-^* and *Il10^-/-^* littermates. DKO and *Il10^-/-^* controls were then separated at weaning based on genotype and each strain rapidly evolved distinct microbiota that was transferrable between hosts. Importantly, horizontal microbial transfer via co-housing DKO mice with *Il10^-/-^* mice attenuated DKO colitis and dysplasia, demonstrating that microbial composition and function may govern disease severity. The microbial profile of co-housed DKO mice more closely mirrored that of *Il10^-/-^* mice. The decreased presence of specific microbial species such as *C. scindens* in singly housed DKO mice may contribute to decreased secondary bile acid production, a critical factor in maintaining intestinal homeostasis^44^. Lynch et al. showed that Turicibacter, decreased in our singly housed DKO mice, could regulate inflammatory pathways by modifying host bile acid profiles and influencing lipid metabolism. Our findings highlight the rapid divergence of the microbiome in littermates with different genetic predispositions that lead to inflammation.

The distinct bile acid metabolomic profiles support a potential role in intestinal inflammation and dysplasia/invasive cancer in DKO mice. Notably, DKO mice had markedly deficient secondary bile acids, which signify disrupted bacterial metabolism. Secondary bile acid supplementation can ameliorate experimental colitis, ^45^ pointing to their therapeutic potential in intestinal inflammation. Fecal LCA concentrations were greater in IBD vs PSC-IBD patients ^46^. The finding of reduced LCA derivatives in DKO mice align with recent insights into their regulatory capacity. LCA and its derivatives, including 3-oxoLCA and isoalloLCA, diminish TH_17_ and promote Treg cell differentiation, that may be anti-inflammatory^33^. We did not identify the bacterial species that perform 3a/b dehydrogenase activity (rate limiting step in 3-oxoLCA metabolism ^34^), perhaps for technical reasons (abundance below 16S detection or unrecognized function of the detected bacteria). Deficient secondary bile acids also occurred in *Mdr2*^-/-^ mice, but this did not promote inflammation; inflammation apparently needed a second hit in DKO mice with loss of IL-10 immunoregulatory function, a finding that mirrors abnormalities seen in PSC and IBD patients ^8,47^.

Reduced luminal LCA derivatives in DKO mice may also promote liver and colon inflammation. Lack of LCA derivatives promotes TH_17_ differentiation ^34^, and IL-17 producing lamina propria TH17 cells and intrahepatic *γδ* cells can intensify colitis and cholestatic liver disease, respectively ^48^ ^26^. Reduced anti-inflammatory Treg cells may be compromised, thereby exacerbating intestinal inflammation and contributing to colitis-associated dysplasia. Bedke et al reported decreased distal colonic Tregs in their DKO model ^39^, although we did not note decreased *foxp3* colonic expression in our mice. Collectively, these insights support microbiota’s dualistic role in modulating PSC-IBD and may guide novel microbiota-centric therapeutic approaches.

One unanticipated result was that DKO mice show less histologic liver fibrosis compared to *Mdr2^-/-^* littermates, despite elevated levels of macrophage chemokines in the DKO mice. PSC patients have increased periportal macrophages ^49^; our findings extend this observation by demonstrating elevated, microbiota-dependent reparative macrophages in DKO mice compared to *Mdr2^-/-^* littermates, likely mediated by increased IL-33 expression that has been shown to induce reparative macrophages ^32^. Notably, beneficial effect is absent in germ-free mice, suggesting a critical microbiota role in this phenotypic switch towards CD206+ macrophages ^50,51^. This phenotypic switch indicates the transition from an inflammatory to a tissue-reparative state, supported by increased MMP activity (MMP-2, -9, -13) and TIMP-1 gene expression. These reparative macrophages produce anti-inflammatory cytokines and mediators, crucial for resolving hepatic fibrosis by remodeling the extracellular matrix and degrading fibrous tissue ^52^.

Although GF DKO mice lack colitis, we observed 100% liver-mediated lethality like *Mdr2^-/-^* mice. GF DKO mice exhibited reduced liver inflammation with no change in liver fibrosis compared to age-matched SPF DKO controls. Microbial-dependent periportal DKO macrophage enrichment is supported by loss of the liver macrophage infiltration and monocyte chemokine expression differences with absent microbiota that were observed between SPF DKO and *Mdr2^-/-^* mice. Microbial reconstitution rescued mortality in most of these mice, underscoring the hepatoprotective role of the DKO microbiota. This highlights the microbiome’s critical influence in modulating liver pathology and its complex role in hepatobiliary fibrosis and inflammation. Microbial protection in DKO mouse suggests promising avenues for identifying specific microbial drivers that could leverage therapeutic interventions

We acknowledge certain limitations of our study. While the DKO model effectively captures several key clinical features of PSC-IBD, it remains a simplified representation of this complex human disease. Intrinsic differences between mouse and human immunology and microbiota may lead to variable disease manifestations and treatment responses. The inability to fully mimic the chronic and progressive nature of human PSC-IBD may overlook long-term disease dynamics. Additionally, the precise mechanisms by which the microbiota influence the gut-liver axis, particularly in the context of observed metabolic alterations, remain to be fully elucidated. Colonizing GF DKO mice with PSC-IBD patients fecal transplants will be critical for translating our findings into potential therapeutic interventions for PSC-IBD.

In conclusion, our study uniquely dissects PSC-IBD pathogenesis by revealing colon-liver-microbiota interactions, uncovering potential avenues for selective microbiota interventions. These findings could guide strategies for microbiome manipulation, potentially altering colitis and hepatobiliary progression.

## Supporting information

supplemental figure legens

## Acknowledgments

The results are supported by NIH grants P01DK094779, National Gnotobiotic Rodent Resource Center (P40OD010995), P30 CGIBD P30DK034987 (to RBS), T32DK07737 and T322A1007273 (to MA), The Crohn’s and Colitis Foundation #2434 (to RBS); VA Merit Award I01BX004033; Research Career Scientist Award (IK6BX004477); VA ShEEP Grant (1 IS1 BX004777); NIH Grants R01 DK104893, R01DK-057543 (to HZ). Bo Liu, DVM, PhD, Jeremy Herzog, Amba Viswanathan and Akihiko Oka, MD, PhD provided technical assistance; Josh Frost, UNC Gnotobiotic Facility Manager Biology for gnotobiotic research support; Josue Hernandez, Fengling Li, MD, Ph.D., DVM for germ-free derivation of *MMdr2*^-/-^ mice; Lisa Holt for laboratory management. The UNC Microbiome Core is supported by Center for Gastrointestinal Biology and Disease (CGIBD P30 DK034987) and the UNC Nutrition Obesity Research Center (NORC P30 DK056350). Raneem Khedraki, PhD for editing support.

## Authorship Contributions

Muyiwa Awoniyi, MD, PhD (Conceptualization: Lead; Data curation: Lead; Formal analysis: Lead; Investigation: Lead;

Methodology: Lead; Project administration: Lead; Validation: Lead; Writing – original draft: Lead; Writing – review & editing: Lead)

Billy Ngo (Data curation: Supporting; Formal analysis: Supporting)

Vik Meadows, PhD (Data curation: Supporting; Investigation: Supporting)

Deniz Coskuner, MD (Data curation: Supporting; Formal analysis: Supporting)

Stephanie A. Montgomery DVM, PhD (Data curation: Supporting; Formal analysis: Supporting)

Morgan Farmer (Data curation: Supporting)

Bo Liu PhD (Data curation: Supporting)

Huiping Zhou PhD (Data curation: Supporting)

Jeffery Roach, PhD (Data curation: Supporting; Formal analysis: Supporting; Methodology: Supporting) Thaddeus Stappenbeck, MD, PhD (Resources: Supporting; Writing – review & editing: Supporting)

Ryan Balfour Sartor, MD (Conceptualization: Lead; Funding acquisition: Lead; Investigation: Lead; Project administration: Lead; Supervision: Lead; Writing – original draft: Lead; Writing – review and editing: Lead)

