## supplemental figures for "Gut Microbial Impact on Colitis and Colitis-Associated Carcinogenesis in a Primary Sclerosing Cholangitis-IBD Model"

### Supplemental Figure 1

**A.**

| Dysplasia Score | Description |
|---|---|
| 0 | Normal glandular architecture |
| 1 | Mild dysplasia: minimal architectural changes (slight focal cellular atypia) |
| 2 | Moderate dysplasia with marked cellular atypia and glandular alterations |
| 3 | Severe dysplasia with significant architectural distortion, marked cellular atypia and glandular crowding |
| 4 | High grade dysplasia exhibiting marked architectural and cytologic abnormalities |
| 5 | Poorly differentiated dysplasia with muscular propria involvement |

**B.**

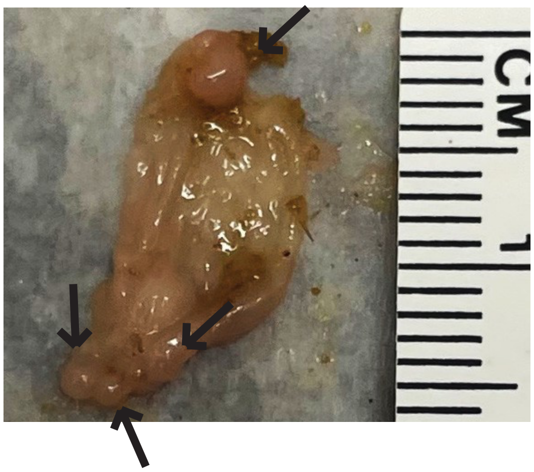


Invasive Ductal Adenocarcinoma (DKO)

**C.**

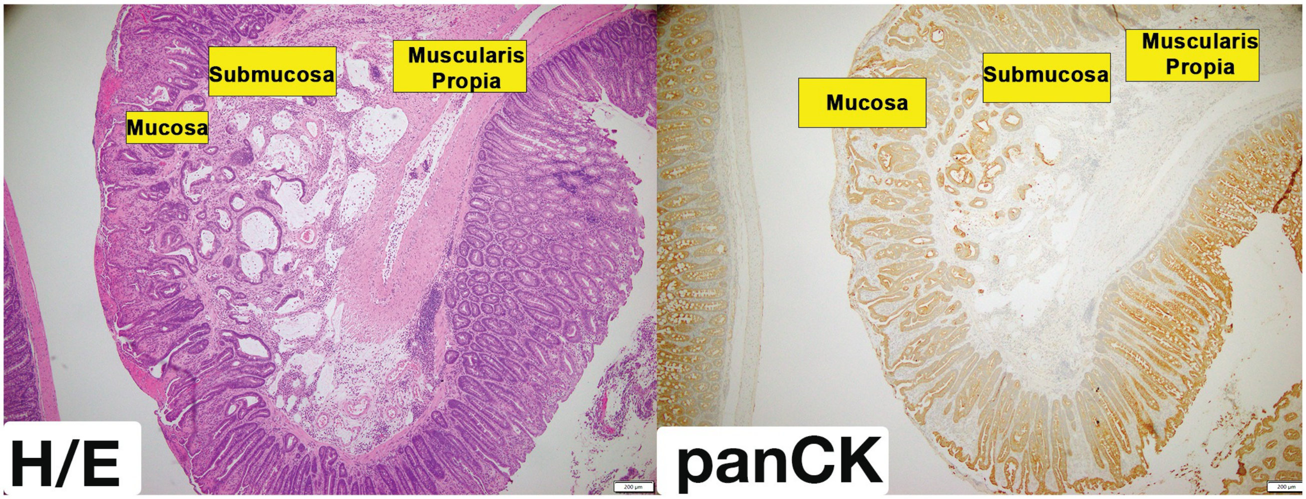


Cecal Cytokine/ Chemokine Map

**D.**

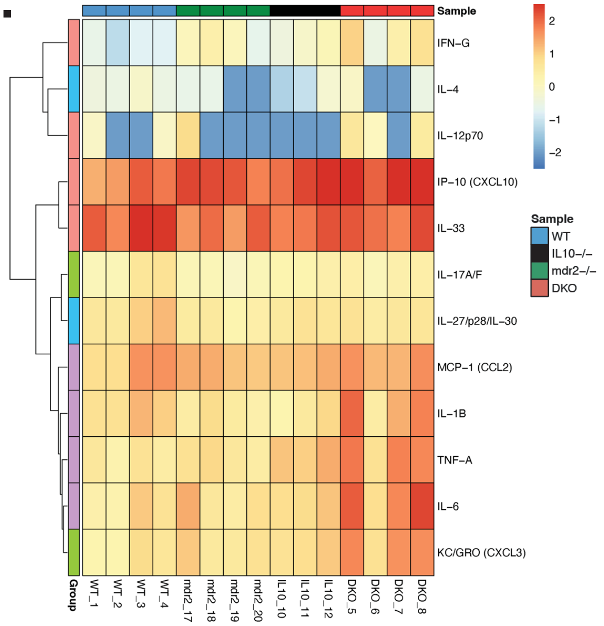


Cytokine/Chemokine (Cecum)

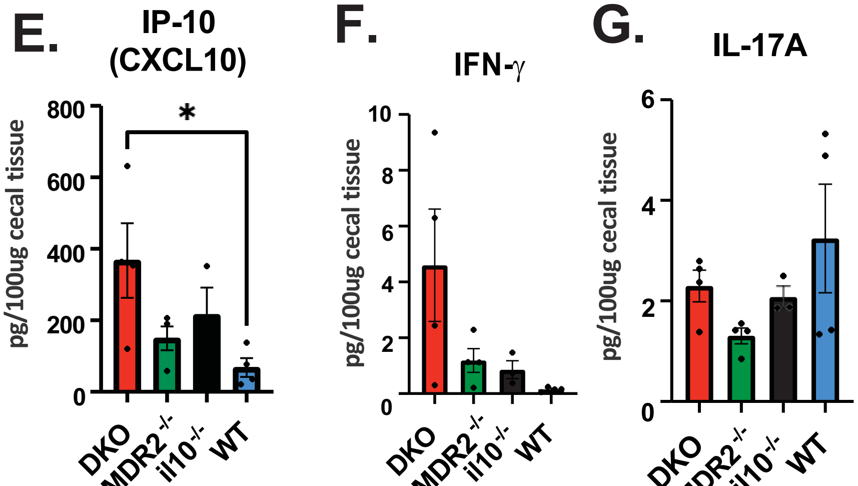

### Supplemental Figure 2

| WT | MDR2-/- | IL-10-/- | DKO |
| --- | --- | --- | --- |
| 6-8 week old | | | |

### Supplemental Figure 3

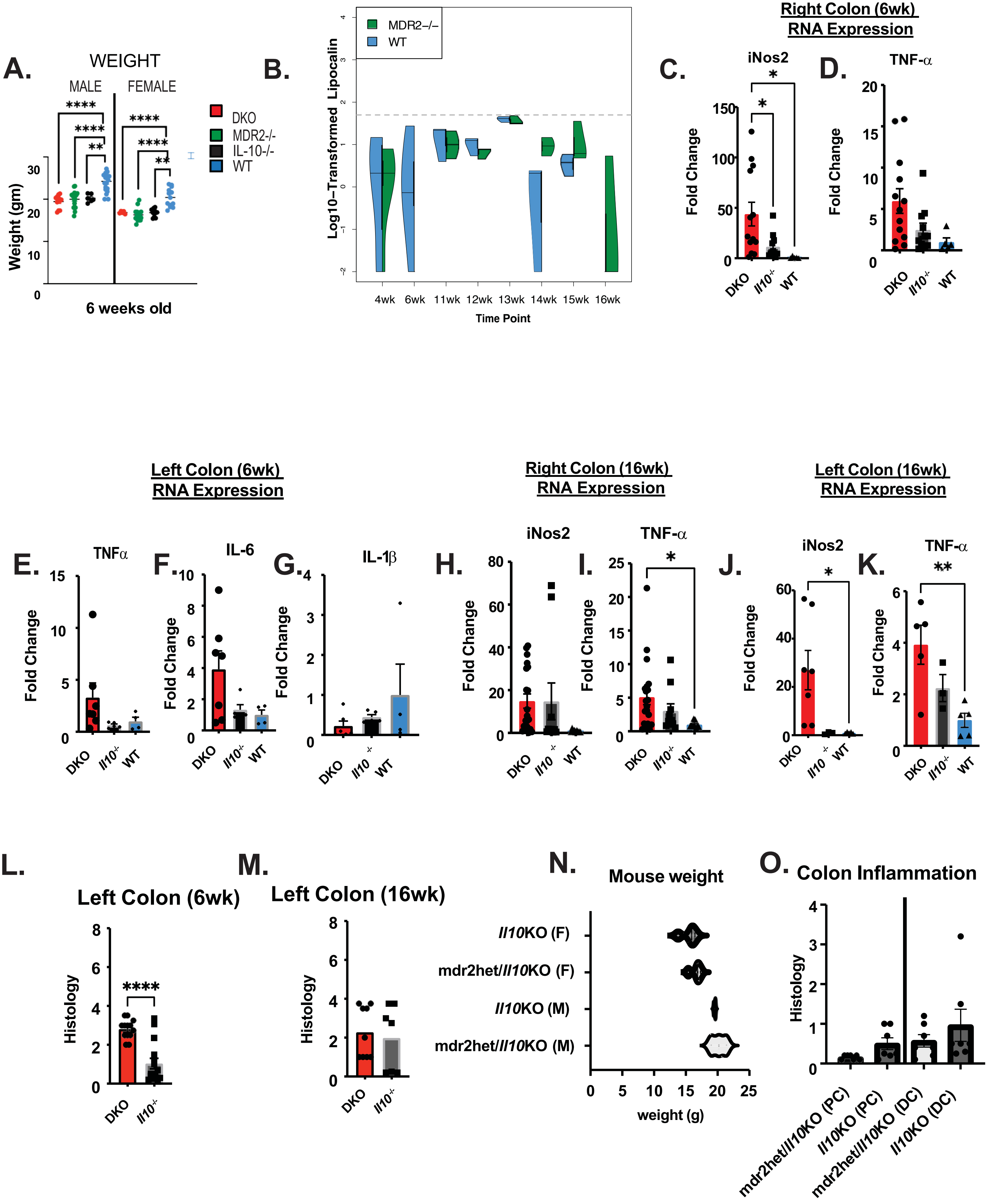

### Supplemental Figure 3

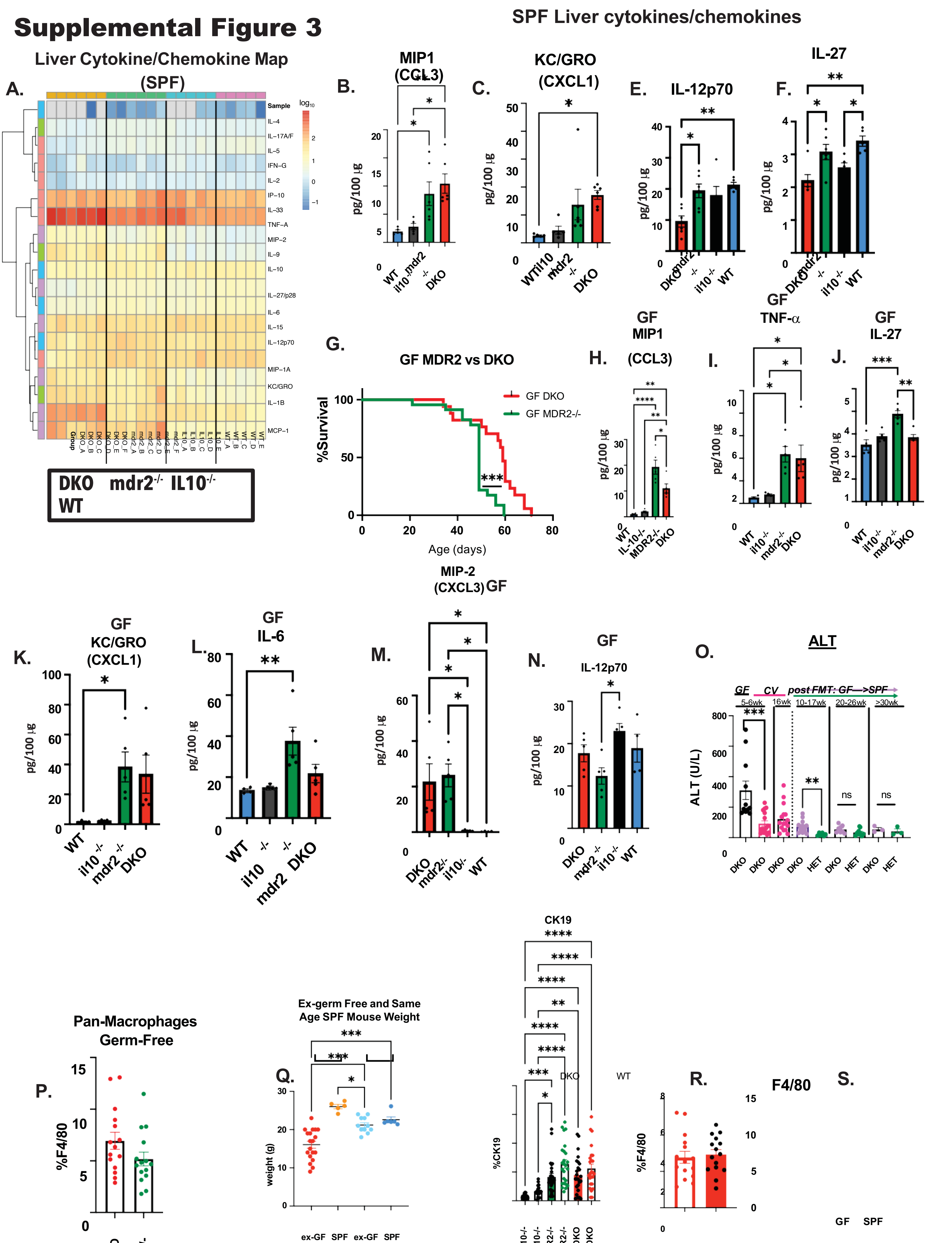

DKO

T.

15

10

5

0

GF SPF

DKO

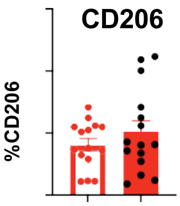

### Supplemental Figure 4

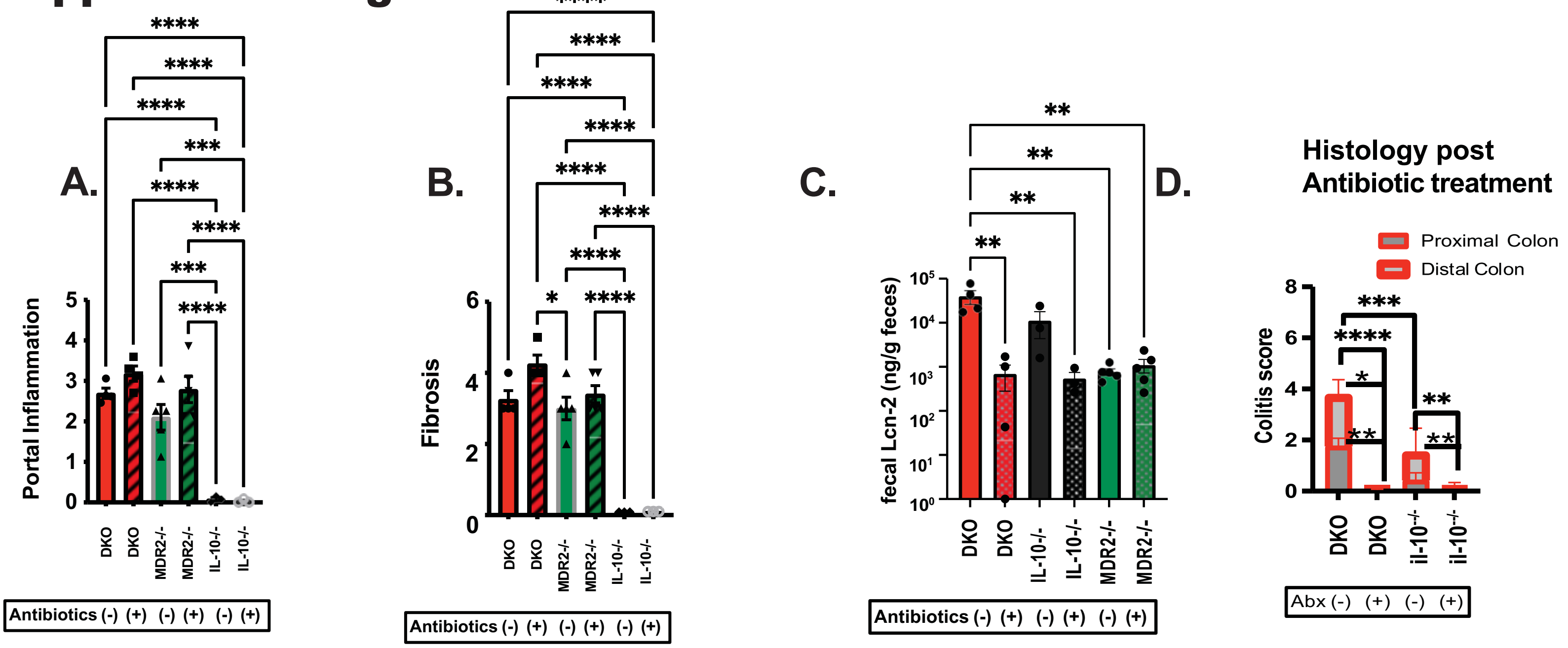

### Supplemental Figure 5

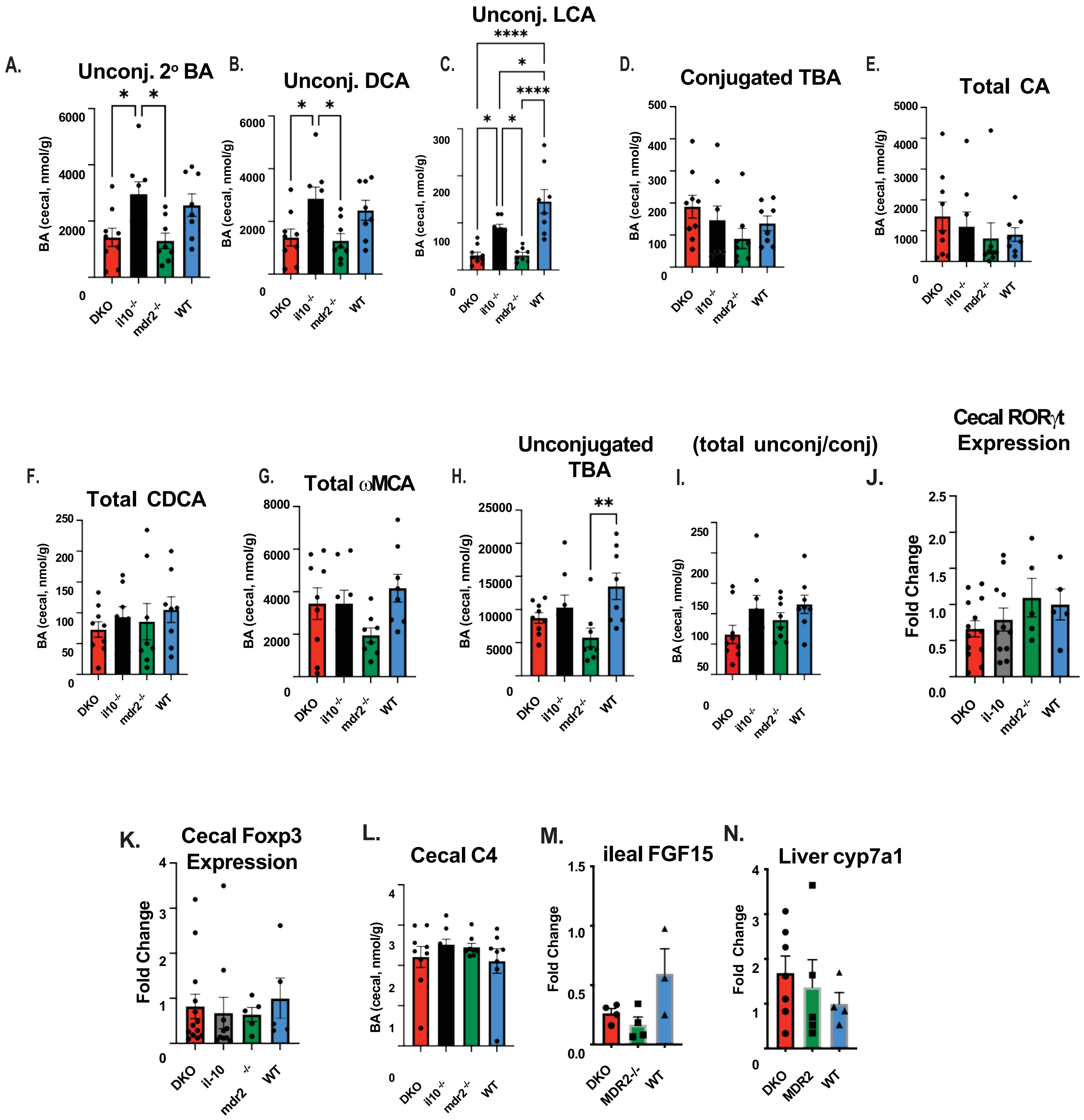

### Supplemental Figure 6

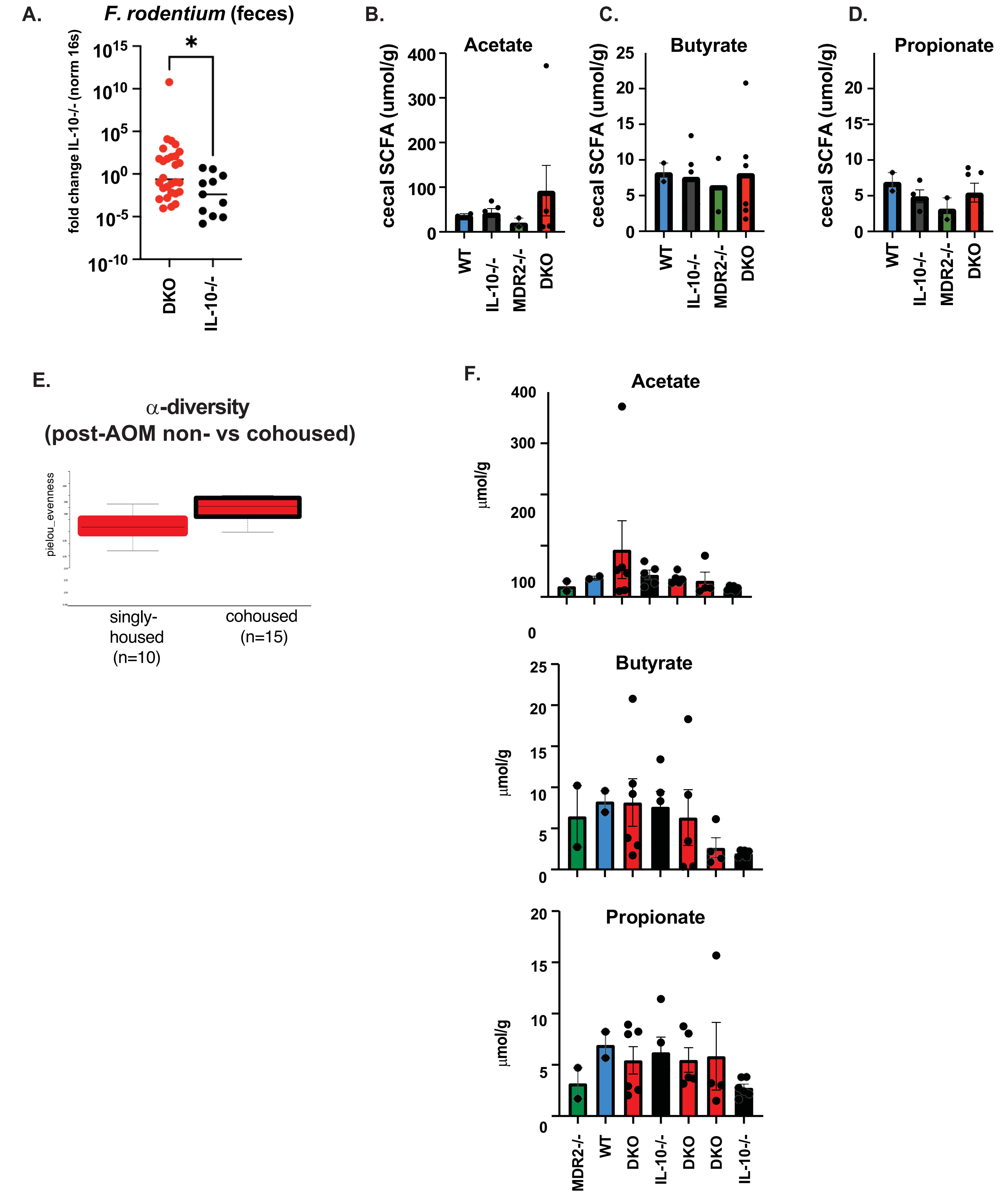

### Supplemental Figure 6

| AOM | (-) | (-) | (-) | (-) | | (+) | (+) | (+) |
|---|---|---|---|---|---|---|---|---|
| co-house | (-) | (-) | (-) | (-) | (-) | | (+) | (+) |

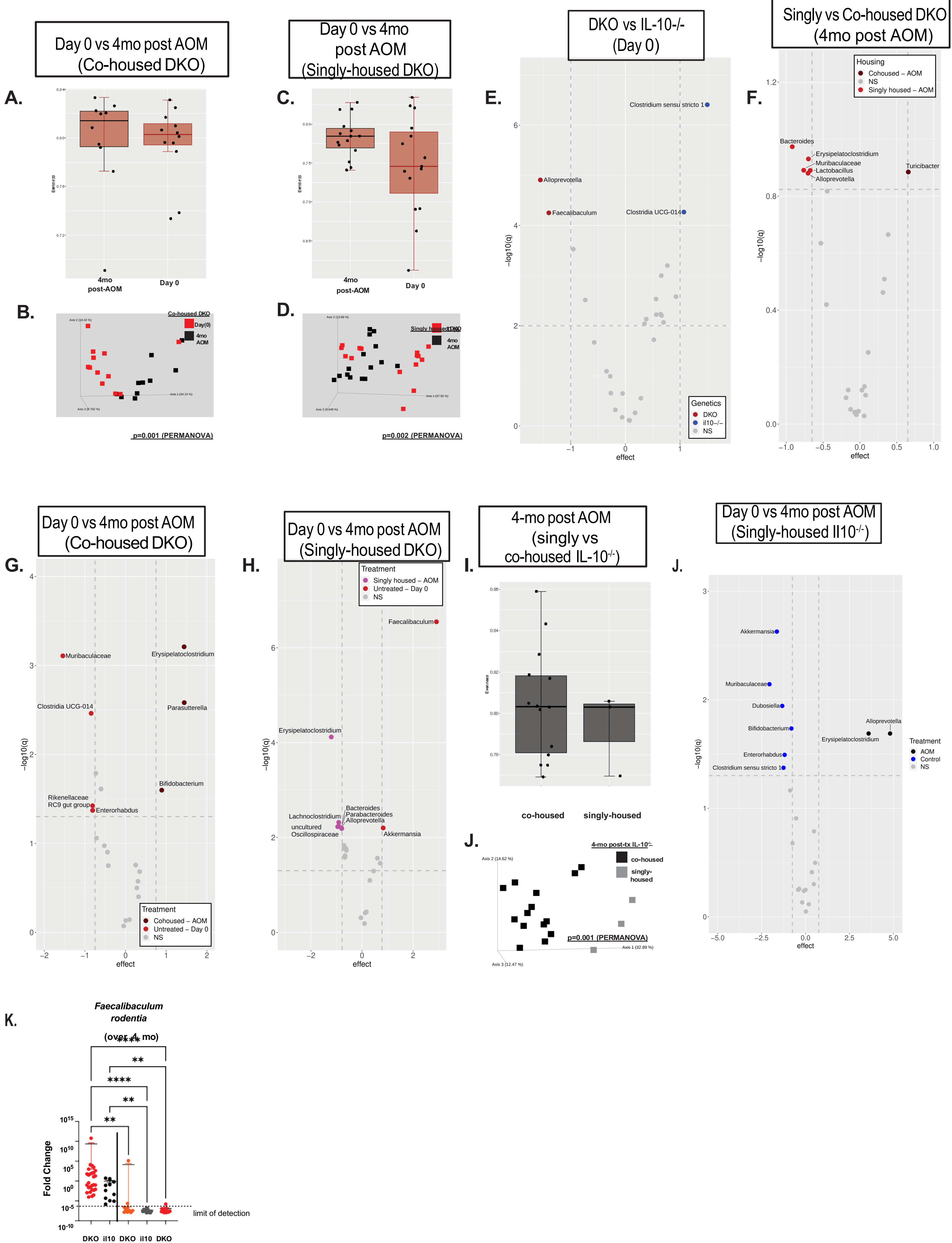
Day 0 vs 4mo post AOM (Co-housed DKO)
A.
4mo post-AOM
Day 0
B.
Co-housed DKO
Day(0)
4mo AOM
p=0.001 (PERMANOVA)
Day 0 vs 4mo post AOM (Singly-housed DKO)
C.
4mo post-AOM
Day 0
D.
Singly housed DKO
4mo AOM
p=0.002 (PERMANOVA)
DKO vs IL-10-/- (Day 0)
E.
Clostridium sensu stricto 1
Alloprevotella
Faecalibaculum
Clostridia UCG-014
Genetics
DKO
il10−/−
NS
−log10(q)
effect
Singly vs Co-housed DKO (4mo post AOM)
F.
Housing
Cohoused – AOM
NS
Singly housed – AOM
Bacteroides
Erysipelatoclostridium
Muribaculaceae
Lactobacillus
Alloprevotella
Turicibacter
−log10(q)
effect
Day 0 vs 4mo post AOM (Co-housed DKO)
G.
Muribaculaceae
Erysipelatoclostridium
Clostridia UCG-014
Parasutterella
Bifidobacterium
Rikenellaceae RC9 gut group
Enterorhabdus
Treatment
Cohoused – AOM
Untreated – Day 0
NS
−log10(q)
effect
Day 0 vs 4mo post AOM (Singly-housed DKO)
H.
Treatment
Singly housed – AOM
Untreated – Day 0
NS
Faecalibaculum
Erysipelatoclostridium
Lachnoclostridium
Bacteroides
Parabacteroides
Alloprevotella
uncultured Oscillospiraceae
Akkermansia
−log10(q)
effect
4-mo post AOM (singly vs co-housed IL-10-/-)
I.
co-housed
singly-housed
J.
4-mo post-tx IL-10-/-
co-housed
singly-housed
p=0.001 (PERMANOVA)
Day 0 vs 4mo post AOM (Singly-housed Il10-/-)
J.
Akkermansia
Muribaculaceae
Dubosiella
Bifidobacterium
Alloprevotella
Erysipelatoclostridium
Enterorhabdus
Clostridium sensu stricto 1
Treatment
AOM
Control
NS
−log10(q)
effect
K.
Faecalibaculum rodentia
(over 4 mo)
****
**
****
**
**
Fold Change
limit of detection
DKO
il10
DKO
il10
DKO

cohoused (-) (-) (+) (+) (-)

**Genetics/Housing**

| Species | DKO vs. $Il10^{-/-}$ | | DKO vs. $Mdr2^{-/-}$ | | $Il10^{-/-}$ vs. $Mdr2^{-/-}$ | |
|---|---|---|---|---|---|---|
| | Effect size | Adj. *p*-value | Effect size | Adj. *p*-value | Effect size | Adj. *p*-value |
| *Escherichia coli* | -0.0167 | 0.8233 | -0.4258 | 0.2740 | -0.4344 | 0.5533 |
| *Bacteroides fragilis* | -0.8613 | 0.0836 | -1.4578 | 0.0317 | -0.5700 | 0.3824 |
| *Helicobacter hepaticus* | -0.9149 | 0.0802 | -0.2582 | 0.3721 | 0.5738 | 0.4513 |
| *Klebsiella oxytoca* | -0.5851 | 0.1979 | -0.0474 | 0.8964 | 0.6207 | 0.4000 |
| *Morganella morganii* | -0.3032 | 0.5300 | -0.1315 | 0.7928 | 0.1324 | 0.8676 |
| *Clostridioides difficile* | 1.4316 | 0.0145 | 0.6889 | 0.1389 | -0.8032 | 0.2617 |
| *Clostridium perfringens** | 1.1585 | 0.0240 | -0.0442 | 0.8815 | -1.4333 | 0.0480 |
| *Salmonella enterica* | -0.5273 | 0.2923 | -0.8791 | 0.0584 | -0.1740 | 0.7426 |
| *Enterococcus faecalis*** | 1.0648 | 0.0310 | 0.1804 | 0.7536 | -0.6762 | 0.1872 |

*Clostridium ramosum*, *Fusobacterium nucleatum*, *Peptostreptococcus anaerobius*, *Campylobacter jejuni*, and *Citrobacter rodentium* not observed.

Species taken from:

https://www.science.org/doi/10.1126/science.abm3233

https://www.tandfonline.com/doi/full/10.1080/19490976.2023.2185028

Notes:

* *Clostridium perfringens* was originally observed in insufficient quantity to be included in the differential abundance analysis; however, with the updated database we now have sufficient reads. It falls in Group 1a. It is unusual in Group 1a in that it also is found in significantly higher relative abundance in $Il10^{-/-}$ versus $Mdr2^{-/-}$. The only other species like this is *Pseudoalteromonas* sp. 3J6. Since *Pseudoalteromonas* sp. 3J6 is a marine bacterium, I suspect it is a misclassification. But is it found in enough quantity to be included in the analysis.

** In the original differential abundance, *Enterococcus faecalis* was close to, but not quite differentially abundant DKO vs. $Il10^{-/-}$ (effect size:0.8201 Adj. *p*-value: 0.0555). But in the updated database, it is. It falls into Group 1a.
