## supplemental methods for "Gut Microbial Impact on Colitis and Colitis-Associated Carcinogenesis in a Primary Sclerosing Cholangitis-IBD Model"

**SUPPLEMENTAL MATERIAL AND METHODS**

**Ethics Statement/Animal Husbandry**

Animal studies were conducted in accordance with NIH guidelines. All experiments were approved and overseen by the Institutional Animal Care and Use Committee (IACUC) at UNC under Protocol IACUC protocol no. 18-266 and Cleveland Clinic Protocol IACUC no. 2281.

**SPF and Germ-Free Double Knockout Mice**

This study utilized specific-pathogen-free (SPF) *Mdr2/Il10* double knockout (DKO), $Il10^{-/-}$, $Mdr2^{-/-}$, and wild-type (WT) C57BL/6 mice were used. Mice were aged 6-45 weeks. The mice used in this study were born from parents that had also been born and raised in SPF conditions over six generations, and they were used at 8 to 16 weeks of age. C57BL/6 background $Mdr2^{-/-}$mice were kindly donated by Yury Popov, M.D., Ph.D. lab for use in this study and housed in an SPF facility for our antibiotic treatment studies. WT and C57BL/6. $Il10^{-/-}$ were originally purchased from Jackson Laboratories (Bar Harbor, ME) and maintained in the Sartor lab for multiple years. SPF DKO mice were generated by crossing $Il10^{-/-}$ and $Mdr2^{-/-}$, mice. DKO, $Il10^{-/-}$, $Mdr2^{-/-}$ and WT C57BL/6 mice were also derived into germ-free (GF) conditions by cesarean sections and breeding colonies were established by the National Gnotobiotic Rodent Resource Center at UNC-Chapel Hill (NIH Grant P40OD010995). $Mdr^{+/-}Il10^{-/-}$ were crossed with each other in order to obtain $Mdr^{-/-}$ and $Il10^{-/-}$ littermates. These mice were separated per genotype at time of weaning with goal genetic strain specific microbiota. GF mice had access to autoclaved water and irradiated Teklad Global Soy Protein-Free Extruded Rodent Diet (2020SX and 5V0F) ad libitum. SPF mice were given the same autoclaved chow. All experiments used age and sex-matched littermate controls, or their immediate descendants. Mice were housed at 37°C and 12-hour light/dark cycles. All necropsies were performed during the light cycle. For the dysplasia progression study, mice were longitudinally monitored from 16 to 45 weeks.

**Assessment of Colitis**

We evaluated the severity of colitis by serial fecal lipocalin-2 ELISA (R&D systems, DY1857) and blinded histologic scoring as described [31].

**Animal Interventional Experiments**

For antibiotic experiments, SPF mice were exposed to ad libitum drinking a mixture of vancomycin (0.5 mg/ml), neomycin (1 mg/ml), and metronidazole cocktail in distilled water with similar approach as referenced in our previous study [53]. For fecal content transplant experiments, we transplanted feces collected from SPF DKO C57BL/6 mice diluted in 10% glycerol in PBS at a ratio of 100 mg of pre-reduced anaerobic feces/ml that was homogenized in a bead beater. The insoluble particles were removed by passing the mixture through a 100 μm filter. Total protein concentration was determined using the BCA assay (Pierce, #23225) to normalize for variations in the number of bacterial cells. The feces mixtures were stored at -80°C until used. We orally gavaged with 200 μl of fecal mixture twice every 3 days during the first week. The mouse survival rate was monitored, and feces collected pre/post fecal transplantation and serum liver function tests were conducted post-mortem.

**Histology**

Colonic and hepatic tissues were fixed, sectioned, and stained using Hematoxylin & Eosin (H&E) and Periodic acid–Schiff (PAS) to assess tissue architecture and dysplasia by the UNC Center for Gastrointestinal Biology & Disease Histology Core. H&E and Sirius red-stained liver and intestinal sections (5 μm) were examined under light microscopy by an animal pathologist masked to all mouse data. The following histopathological features were assessed: mean score from a minimum of 12 portal systems over at least two lobes assessing for colon and liver inflammation as well as liver fibrosis. Neoplastic lesions were scored by a mouse pathologist using a system based upon a 2003 consensus report on mouse models of intestinal cancer.

**Co-housing DKO and Il10$^{-/-}$ Model**

For the co-housing experiment, DKO and *Il10*$^{-/-}$ mice were born and raised in our SPF facility (age 6-8 weeks, singly or co-housed 2-5 mice per cage) for the immediate initiation of our colitis/colorectal cancer experiment. At weaning, we then separated genotype-specific strains that led to the divergence of the microbiome composition described in Fig. Mice received 4 weekly i.p. injections of AOM (10 mg/kg, Sigma-Aldrich, A5486) or PBS. Mice underwent necropsy at the 4 months following the initial injection, and stool and tissues were collected, and colons were examined macroscopically for tumors and fixed in 10% formalin for paraffin embedding and histology.

**Immunohistochemical Staining and Quantification of Hepatic Macrophage Markers**

Liver tissue sections (5 μm) following deparaffinization and rehydration underwent heat-induced antigen retrieval and endogenous peroxidase blocking. Sections were washed and blocked for non-specific binding with 2.5% horse serum and incubated overnight with primary antibodies against F4/80 (a pan-macrophage marker), CD206 (a marker for anti-inflammatory, reparative macrophages), and IRF5 (a marker for pro-inflammatory macrophages). After thorough washing, immunoreactivity was performed with ImmPress horseradish peroxidase detection kits. Sections were exposed to an HRP-conjugated secondary antibody, thoroughly washed, and the signal was developed using a DAB substrate for deposition of brown precipitate at the antigen site. Hematoxylin was used for counterstaining to visualize the nuclei.

Stained slides were scanned with a high-resolution digital scanner and three digital non-overlapping images were used for semi-quantification of positive staining. Quantitative analysis was performed using ImageScope through the selection of positively stained areas over the whole field area for each image. All quantifications were performed in a blinded manner to avoid bias. We compared macrophage densities between experimental groups.

**Tissue Cytokine/Chemokine Assessment**

Frozen liver and cecal samples were used to measure cytokine and chemokine concentrations using the mouse V-PLEX Proinflammatory Panel 1 and Cytokine Panel assay (Mesoscale Devices). Prior to assay determination, 50–150 mg samples were homogenized in 1 mL of lysis buffer (150 mM NaCl, 20 mM Tris, 1 mM EGTA, 1% Triton X-100, protease inhibitor) for 4 min at 20 Hz using a tissue homogenizer. Homogenized samples were then centrifuged at 14,000 x g for 10 min to remove debris and appropriately diluted according to total proteins present in corresponding supernatants, as determined by the Pierce BCA Protein Assay Kit (Thermo Scientific, Product No. 23225). Acquired Meso Scale Discovery (MSD, V-Plex Cytokine Panel 1) data for each sample were normalized to its total protein concentration prior to statistical analysis.

Quantitative PCR was employed for validation of chemokine/cytokine/MMP's mRNA expression analysis. Consistent snap-frozen right hepatic lobes were stored in RNAlater (Thermo Fisher, AM7020) at -80°C. A small portion was removed from the storage solution, and the tissue was homogenized. RNA was isolated via the RNeasy Mini Kit (QIAGEN, #74104) according to the manufacturer's recommendations. cDNA was obtained by reverse transcription of 250 ng of total RNA using iScript cDNA Synthesis Kit (Biorad, 1708890) according to the instructions of the manufacturer. Relative mRNA transcript levels were quantified using TaqMan Universal PCR Master Mix (Applied Biosystems) applying the TaqMan methodology or iTaq Universal SYBR Green Supermix (Biorad) for SYBR green probes. The housekeeping gene 18s was amplified in a parallel reaction for normalization. Primers for the following genes were used: Monocyte Chemoattractant Protein-1 (MCP-1),

Lipocalin-2 (Lcn-2), Collagen 1a1 (Col1a1), Tissue inhibitor of metalloproteinases (Timp)-1, inducible nitric oxide (iNos)-2, Interleukin-(IL)-6, Tumor Necrosis Factor (TNF)-α, interferon gamma (IFN-γ), Rorgt, and FOXP3. Primer sequences are detailed in previous study [53]. All TaqMan probes are positioned on exon–exon boundaries of corresponding genes to exclude co-amplification of genomic DNA. The relative expression of each sample was first normalized to the expression of the reference gene 18s, and then normalized to the average expression in samples from Mdr2-/- mice. The data were analyzed according to the 2-ΔΔCT method. Table 1

**Matrix Metalloproteinase Activity Assay**

Matrix metalloproteinase (MMP) activity was measured using the MMP activity assay kit (Abcam; ab112146), in accordance with the manufacturer's instructions. Flash-frozen liver from the median lobe tissue is first homogenized in RIPA buffer. The homogenate was centrifuged at 15,000xg, saving supernatant for use in the lysate MMP activity assay. Protein concentration was determined using a BCA assay. The concentrated samples were adjusted to the same volume. The activities of the MMPs were measured with a fluorescence plate at 45 minutes following the terminal step with the reader at excitation/emission wavelengths of 490/525 nm.

**Biochemical Assessment of Fibrosis**

Hydroxyproline (HYP) was determined biochemically as previously described. Briefly, two snap-frozen liver pieces from the left and right liver lobe (100-150 mg additive total) were hydrolyzed in 5 ml 6 N HCl at 110°C for 16 h. Supernatants were transferred to a 96-well plate, and wells were allowed to evaporate dry. Hydroxyproline content was determined by colorimetric analysis (catalog # MAK008, Sigma Aldrich, St. Louis, MO, USA) and expressed as μg hydroxyproline/mg liver.

**Serum Collection and Liver Function Tests**

Blood was collected by cardiac puncture at the time of mouse necropsy, and serum was separated using BD serum separator additive microtainer tubes centrifuged at 4000 x g for 15 minutes. The recovered serum was stored at -80ºC. Stored serum was sent to the UNC Animal Histopathology and Lab Medicine Core per protocol for liver function studies including Alanine aminotransferase, Alkaline Phosphatase, and Total Bilirubin.

**Fecal Sample Collection and DNA Isolation**

Fecal samples were collected aseptically from live mice and immediately snap-frozen, then stored at -80°C until further processing. DNA isolation commenced by incubating fecal material in Lysing Matrix E tubes (MP Biomedicals) with a lysis buffer containing 200 mM NaCl, 100 mM Tris-HCl (pH 8.0), 20 mM EDTA, SDS, and proteinase K (QIAGEN). The mixture was then subjected to mechanical disruption using a bead beater homogenizer at 4°C for 3 minutes. Following homogenization, phenol:chloroform:isoamyl alcohol (25:24:1) (Invitrogen) was added, and samples were centrifuged at 8000 rpm for 3 minutes at 4°C. The aqueous phase was subsequently treated with an equal volume of phenol:chloroform:isoamyl alcohol and incubated for 10 minutes at room temperature, followed by centrifugation at 13,000 rpm for 5 minutes at 4°C. DNA was precipitated by adding isopropanol and 3M sodium acetate (pH 5.2) and incubated at -20°C for 15 hours. The precipitate was pelleted by centrifugation at 13,000 rpm for 20 minutes at 4°C, washed with cold ethanol, and re-suspended in TE buffer. DNA purification was finalized using the DNeasy Blood and Tissue Kit (QIAGEN) as per the manufacturer's instructions.

**16S rRNA Gene Sequencing**

The 16S rRNA gene sequencing was performed as previously described [53].

**Whole Genome Sequencing (WGS)**

For whole genome sequencing, genomic DNA was extracted from liver tissue using a similar protocol as described above, ensuring high molecular weight DNA was obtained. The DNA was then quantified, and its quality assessed using a Qubit fluorometer and agarose gel electrophoresis. Library preparation for WGS was performed using the Nextera DNA Flex Library Prep Kit (Illumina), following the manufacturer's instructions. Libraries were quantified, normalized, and pooled for sequencing. WGS was conducted on an Illumina NovaSeq 6000 system to achieve a depth of 30x coverage per genome. Sequencing output from the Illumina platform was converted to fastq format and demultiplexed using Illumina BclConvert. Quality control of the demultiplexed sequencing reads was verified by FastQC. Adapters were trimmed using Trim Galore. The resulting paired-end reads were submitted to Kraken2 for taxonomic classification. An estimate of taxonomic composition including host was produced from these results using Bracken 2.5. All reads classified as host were eliminated. Diversity analysis: alpha diversity as measured by Evenness index and beta diversity as measured by Bray-Curtis dissimilarity was performed using QIIME2. Differential abundance was estimated with

ALDEx2 and ANCOM-BC on species with a minimum abundance of 5000 reads and minimum prevalence of 20%. Differential abundant species (transpose) principal coordinate analysis was performed on all species found to be differentially abundant at the $q<0.05$ level by ALDEx2.

**Microbial Analysis**

Sequencing output from the Illumina MiSeq platform was converted to fastq format and demultiplexed using Illumina BCL Convert 3.8.2-12. The resulting paired-end reads were processed with the QIIME 2 2022-2 wrapper for DADA2 including merging paired ends, quality filtering, error correction, and chimera detection. Amplicon sequencing units from DADA2 were assigned taxonomic identifiers with respect to the Silva database, their sequences were aligned using maFFT in QIIME 2, and a phylogenetic tree was built with FastTree in QIIME 2. Alpha diversity with respect to Faith index and Evenness index metrics was estimated using QIIME 2 at a rarefaction depth of 5,000 sequences per subsample. Beta diversity estimates were calculated within QIIME 2 using weighted Unifrac distances and Bray Curtis distance between samples at a subsampling depth of 10,000. Results were summarized and visualized by principal coordinate analysis as implemented in QIIME 2. Differential abundance of genera was visualized using the QIIME 2 ANCOM plug-in.

**Targeted Bile Acid Metabolomic Analysis**

All analyses of bile acid metabolites in cecal contents were performed at Huiping Zhou's, PhD, Chemistry and Analytic Core at Virginia Commonwealth University. A UHPLC/Q-TOF method was performed as previously described. Vitamers were analyzed by liquid chromatography-tandem mass spectroscopy (LCMS/MS) whereas short-chain fatty acids (SCFA) were analyzed by gas chromatography-tandem mass spectroscopy (GC-MS/MS) at the CCF Metabolomics Core Facility.

**Statistical Analysis**

Statistical significance was assessed using ANOVA for multiple group comparisons and Student's t-test for pairwise analyses. Correlation coefficients were calculated to assess associations between variables. Survival rates were analyzed using Kaplan-Meier estimates and the Log-rank test. Principal coordinates analysis (PCoA) was utilized to display variations in cytokine, chemokine, and microbial profiles among genotypes. PERMANOVA was employed for multivariate statistical analyses of beta diversity of microbiota data. All data are presented as mean ± SEM, with significance levels and specific p-values including multiple test adjusted p-values (q-values) where appropriate, indicated throughout.

Statistical significance of differentially abundant genera was estimated with ALDEx2 and ANCOM-BC. Statistical significance of pairwise differences in alpha diversity and pairwise distances in beta diversity were estimated using the QIIME 2 q2-longitudinal plug-in. Estimates of paired differential abundance were calculated using negative binomial and zero-inflated mixed models (NBZIMM) as implemented.
