## supplemental figure legens for "Gut Microbial Impact on Colitis and Colitis-Associated Carcinogenesis in a Primary Sclerosing Cholangitis-IBD Model"

**SUPPLEMENTARY FIGURE LEGENDS**

**Supplemental Figure 1: Dysplasia Scoring and Cecal Cytokine/Chemokine Profiles in DKO and Control Mice**

(A) Table summarizing the dysplasia scoring system used to evaluate colonic tissue. (B) Image of representative cecal polyps, marked by arrows 40-week-old DKO mouse (largest 30mm). (C) Sample photomicrograph invasive colonic adenocarcinoma. Left panel: H&E staining showing tissue architecture at 40x magnification. Right panel: panCK staining at 40x magnification highlighting epithelial cells. (D) Heatmap showing the cecal cytokine/chemokine profiles in DKO, *Mdr2*$^{-/-}$, *Il10*$^{-/-}$, and WT mice. Bar graphs depicting the levels of IP-10 (CXCL10), IFN-γ, and IL-17A (E-G) in cecal tissue of DKO, *Mdr2*$^{-/-}$, *Il10*$^{-/-}$, and WT mice. Data are presented as mean ± SEM, with statistical significance indicated by *P < 0.05.

**Supplemental Figure 2: Inflammation and RNA Expression in DKO and Control Mice**

(A) Weight measurements of male and female DKO, *Mdr2*$^{-/-}$, *Il10*$^{-/-}$, and WT mice at 6 weeks old. (B) Longitudinal log10-transformed lipocalin-2 levels in fecal samples from *Mdr2*$^{-/-}$ and WT mice over 16 weeks. RNA expression levels of iNOS2 (C) and TNF-α (D) in the right colon of 6-week-old DKO, *Il10*$^{-/-}$, and WT mice. RNA expression levels of TNF-α (E), IL-6 (F), and IL-1β (G) in the left colon of 6-week-old DKO, *Il10*$^{-/-}$, and WT mice. RNA expression levels of *inos2* (H), TNF-α (I), *inos2* (J), and TNF-α (K) in the right and left colon of 16-week-old DKO, *Il10*$^{-/-}$, and WT mice. Histological scores of the left colon in 6-week-old (L) and 16-week-old (M) DKO and *Il10*$^{-/-}$ mice. (N) Mouse weights and (O) colitis scoring of 6 week of male and female littermates *Il10*$^{-/-}$ and *Mdr2*$^{+/-}$/ *Il10*$^{-/-}$ mice. Data are presented as mean ± SEM, with statistical significance indicated by **P < 0.01, ***P < 0.001, ****P < 0.0001.

**Supplemental Figure 3: Liver Cytokine/Chemokine Profiles and Fibrosis in DKO and Control Mice**

**(**A) Heatmap showing liver cytokine/chemokine profiles in SPF DKO, *Mdr2*$^{-/-}$, *Il10*$^{-/-}$, and WT mice. (B-F) Bar graphs depicting levels of MIP1 (CCL3), KC/GRO (CXCL1), IL-12p70, and IL-27 in liver tissue of SPF DKO, *Mdr2*$^{-/-}$, *Il10*$^{-/-}$ and WT mice. (G) Survival curve comparing GF DKO and GF *Mdr2*$^{-/-}$ mice. (H-N) Liver protein levels of MIP1 (CCL3), TNF-α, IL-27, KC/GRO (CXCL1), IL-6, MIP-2 (CXCL3), and IL-12p70 in liver tissue of GF DKO, *Mdr2*$^{-/-}$, *Il10*$^{-/-}$ and WT mice. (O) Serum alanine transaminase (ALT) levels in DKO, *Il10*$^{-/-}$, and *Mdr2*$^{-/-}$ mice post-FMT, and ex-GF mice over time. (P) Percentage of F4/80 positive macrophages in liver tissue of GF

DKO and *Mdr2*$^{-/-}$ mice. (Q) Comparison of body weight between ex-GF and SPF DKO and WT mice. (R) Quantification of IHC stained CK19 positive, F4/80 and CD206 positive liver tissue of GF and ex-GF DKO and *Mdr2*$^{-/-}$ and *Il10*$^{-/-}$ mice.

**Supplemental Figure 4: Impact of Antibiotics on Inflammation and Fibrosis**

(A-B) Portal inflammation and liver fibrosis scores in DKO, *Il10*$^{-/-}$, and *Mdr2*$^{-/-}$ mice with and without antibiotic treatment. (C-D) Fecal lipocalin-2 levels and colon inflammation score in DKO, *Il10*$^{-/-}$, and *Mdr2*$^{-/-}$ mice with and without antibiotic treatment. N=3-5mice /group.

**Supplemental Figure 5: Bile Acid Profiles in DKO and Control Mice**

Histogram showing levels of various cecal bile acids from DKO, *Il10*$^{-/-}$, *Mdr2*$^{-/-}$, and WT mice. Measured bile acids include (A) unconjugated secondary bile acids, (B) unconjugated deoxycholic acid (DCA), (C) unconjugated lithocholic acid (LCA), (D) conjugated total bile acids (TBA), (E) total cholic acid (CA), (F) total chenodeoxycholic acid (CDCA), (G) total w-muricholic acid (MCA), (H) total unconjugated bile acids and (I) unconjugated/conjugated ratio. (J-K) Fold change in cecal RORγt and Foxp3 expression in DKO, *Il10*$^{-/-}$, *Mdr2*$^{-/-}$, and WT mice. (L) Fecal C4 and fold change of (M) ileal FGF15 and (N) liver cyp7a1expression in DKO, *Mdr2*$^{-/-}$, and WT mice. N-6-8 mice/group.

**Supplemental Figure 6: Microbial Profiles and Short-Chain Fatty Acids (SCFA) in DKO and Control Mice**

(A)Fold change in *Faecalibaculum rodentium* levels in fecal samples from 8-week-old DKO and *Il10*$^{-/-}$ mice. Baseline levels of SCFAs, including (B) acetate, (C) butyrate and (D) propionate, in cecal contents of DKO, *Il10*$^{-/-}$, *Mdr2*$^{-/-}$, and WT mice.  (E) Alpha diversity (Pielou’s evenness) post-AOM treatment in singly housed versus co-housed DKO mice. (F) Levels of cecal SCFAs of singly housed and co-housed DKO and *Il10*$^{-/-}$ mice post-AOM treatment relative to *Mdr2*$^{-/-}$ and WT control mice.

**Supplemental Figure 7: Microbial Community Analysis and Fecalibaculum rodentium Levels in DKO Mice Post AOM Treatment**

(A) Alpha diversity (Pielou's evenness) in co-housed DKO mice compared between day 0 and 4 months post-AOM treatment. Box plots indicate no significant change in microbial evenness over time. (B) PCoA based on Bray-Curtis dissimilarity showing microbial community changes in co-housed DKO mice between day 0 and 4 months post-AOM treatment. PERMANOVA p=0.001 indicates significant differences in microbial composition over time. (C) Alpha diversity (Pielou's evenness) in singly-housed DKO mice compared between day 0 and 4 months post-AOM treatment. Box plots indicate no significant change in microbial evenness over time. (D) PCoA based on Bray-Curtis dissimilarity showing microbial community changes in singly-housed DKO mice between day 0 and 4 months post-AOM treatment. PERMANOVA p=0.002 indicates significant differences in microbial composition over time.

(E-J) Differentially abundant bacterial taxa in various comparisons:(E) DKO vs. *Il10*$^{-/-}$ at day 0 showing specific bacterial taxa significantly different between the genotypes. (F) Singly-housed vs. co-housed DKO mice 4 months post-AOM treatment showing specific bacterial taxa significantly different between the housing conditions. (G) Day 0 vs. 4 months post-AOM in co-housed DKO mice showing specific bacterial taxa significantly different over time. (H) Day 0 vs. 4 months post-AOM in singly-housed DKO mice showing specific bacterial taxa significantly different over time. (I-K) Alpha and beta diversity and differential abundance of Day 0 vs. 4 months post-AOM in singly-housed vs co-housed *Il10*$^{-/-}$ mice. (L) Fold change in *Faecalibaculum rodentium* levels in fecal samples from co-housed and singly-housed DKO and *Il10*$^{-/-}$ mice over 4 months. Data are presented as mean ± SEM, with statistical significance indicated by **P < 0.01, ***P < 0.001, ****P < 0.0001.

**Supplemental Table 1: Comparative Analysis of Genotoxic bacterial species**

Differential abundance of various bacterial species across three different groups based on WGS data: DKO vs. Il10-/-, DKO vs. Mdr2-/-, and Il10-/- vs. Mdr2-/-. For each species, the effect size and adjusted p-values are provided. Species such as *Clostridium ramosum*, *Fusobacterium nucleatum*, *Peptostreptococcus anaerobius*, *Campylobacter jejuni*, and *Citrobacter rodentium* were not observed in the analysis. Notably, *Clostridium perfringens* and *Enterococcus faecalis* exhibited differential abundance in updated database analyses.
